# Task demands dynamically structure feature selection, routing, and integration in the human brain

**DOI:** 10.64898/2026.04.15.718655

**Authors:** Yuening Yan, Jiayu Zhan, Yaocong Duan, Oliver G. B. Garrod, Robin A. A. Ince, Chen Zhou, Rachael E. Jack, Philippe G. Schyns

## Abstract

Flexible visual categorization requires selecting task-relevant information from complex stimuli and transforming it through structured computations. Yet how task demands flexibly control these computations in the human brain remains unresolved. Dynamic faces provide a powerful test case, conveying stable identity through 3D shape and transient emotional expression through 4D movement. Using generative 4D face modelling, participant-specific MEG (N = 24), and information-theoretic analyses, we show that the brain implements a structured processing architecture. Task relevance first gates feature representations in occipital cortex, selectively sustaining relevant features while suppressing irrelevant ones. These features are then routed along pathway-specific channels—3D identity features to the ventral pathway and 4D emotion features to the lateral pathway. Finally, these inputs converge in temporal cortex, where identity and emotion features are synergistically integrated for learned identities. These results show how task demands dynamically structure feature selection, gating, routing, and integration, thereby constraining the computations that support flexible categorization.

---

The human brain extracts multiple sources of information from dynamic faces^1–12^, including stable identity^13–15^ and transient emotional expression^16–20^, to guide behavior. Faces therefore provide a powerful model system because they simultaneously convey invariant structural features (e.g., 3D identity cues) and dynamic movement-based features (e.g., 4D expressive features–Action Units). A central question in cognitive neuroscience is how these distinct feature classes are selectively represented, communicated, and integrated to support behavior under changing cognitive task demands. Crucially, identifying where and when specific features are maintained or attenuated, communicated, and integrated does more than describe neural dynamics: it reveals the information-processing architecture underlying cognition, thereby constraining the computations applied to these features and the principles governing flexible categorization.

Anatomical and functional studies have established partially dissociable processing pathways: a ventral occipito-temporal stream supporting identity representations^21–23^ and a lateral occipito-temporal stream supporting expression and socially dynamic information^24–30^. While this division of labour is well documented^31,32^, how task goals dynamically organize feature processing across these pathways remains unclear. In particular, it is unknown whether early task-dependent gating determines subsequent routing and integration of identity and expression information. Without resolving these feature-specific information-processing dynamics, apparent specialization remains compatible with multiple computational architectures, leaving the underlying algorithmic organization underdetermined.

Recent work shows that task-irrelevant facial features can be attenuated at early visual stages, limiting their propagation to higher-level regions^33,34^. However, whether such early attenuation governs downstream pathway-specific routing and cross-stream integration has not been directly tested. Addressing this requires methods that resolve neural dynamics at millisecond precision and track feature-specific information transmission across cortical regions. Establishing whether feature maintenance, routing, and integration unfold in a structured temporal sequence enables direct tests of dependency relations that define and constrain the computations transforming facial features into behavior.

Here we combine generative 4D face modelling^35^, participant-specific magnetoencephalography (MEG), information-theoretic feature analyses^36^ and directed feature information (DFI)^37^ to quantify when and where identity (F_Id_) and emotion (F_Emo_) features are represented and communicated during identity, emotion, and dual-task categorization. As generative modelling explicitly parametrizes stimulus features, and DFI quantifies their directed transmission, this framework allows us to move beyond response similarity and directly interrogate feature-level computations in brain networks at Marr’s algorithmic level ^38^. This approach enables us to test a structured sequence of dependencies (see Figure 1): (i) Stage 1: whether task-irrelevant features are attenuated and task-relevant features are maintained at early occipital (OCC) stages; (ii) Stage 2: whether task-relevant features exhibit pathway-biased directional transmission consistent with ventral and lateral specialization; and (iii) Stage 3: whether integration in temporal cortex (TC) emerges only when both feature classes are maintained and communicated directionally^39,40^.

**Figure 1.**
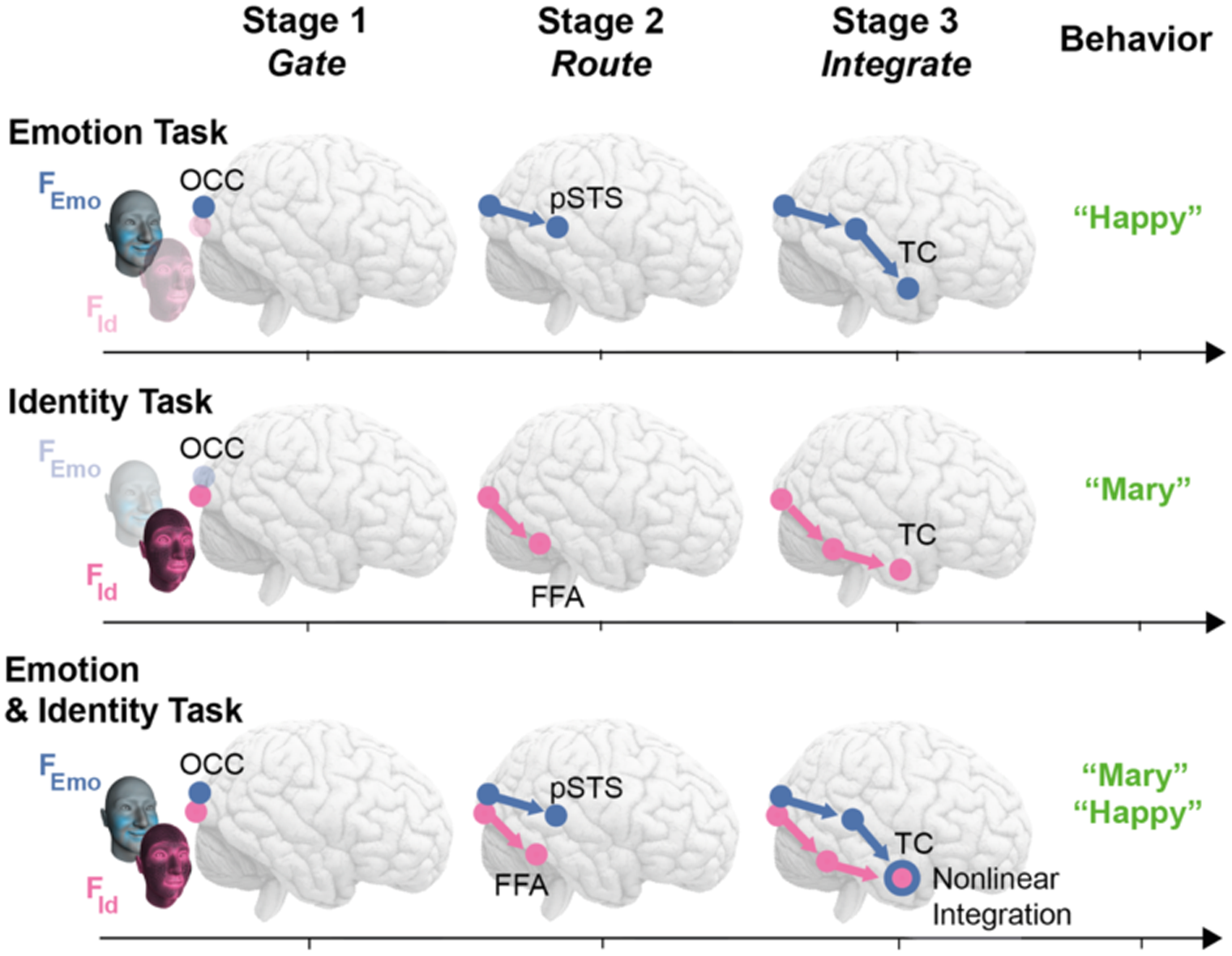
Structured sequence of feature gating, routing, and integration underlying behavior in the ventral and lateral pathways. **Stage 1–Feature gating:** Does OCC maintain task-relevant features and attenuate task-irrelevant features (shaded)? **Stage 2–Directional routing:** Do ventral and lateral pathways directionally communicate task-relevant features (ventral for F_Id_, lateral for F_Emo_)? **Stage 3—Conditional integration:** Does TC integrate F_Id_ and F_Emo_ only when both are task-relevant, maintained, and directionally communicated?

If these stages form a strict dependency structure, then early feature maintenance must constrain downstream routing, and routing must condition integration, thereby limiting the class of computational architectures capable of implementing flexible face categorization.

By quantifying feature-specific representation, directional information flow, and synergistic (nonlinear) integration within a unified framework, this study establishes empirical constraints on how task demands govern the maintenance, attenuation, routing, and integration of facial information to support behavior. More broadly, tracing the spatiotemporal trajectory of stimulus features through the brain reveals the information-processing architecture underlying cognition and constrains the computations that enables flexible visual categorization.

## Results

Using a generative model of the human face^35^, we orthogonally manipulated stable (3D shape) and transient (Action Unit, AU ^41^) facial features. This orthogonal manipulation ensured that identity-defining (F_Id_) and emotion-defining (F_Emo_) features varied independently, allowing their neural representation and communication to be dissociated. We generated twelve random face identities (Figure 2A) and six emotion models^20^–happy, surprise, fear, disgust, anger, and sad (Figure 2B)–each defined by a distinct combination of AUs (see *Methods, Experiment, Stimuli*). Before MEG scanning, participants learned to name six randomly assigned identities (“Known”), allowing us to test whether minimal semantic associations influence downstream feature integration. To ensure equal proficiency across tasks, all participants were also trained to classify the six emotions to 100% accuracy (see *Methods, Experiment, Training Procedure*).

**Figure 2.**
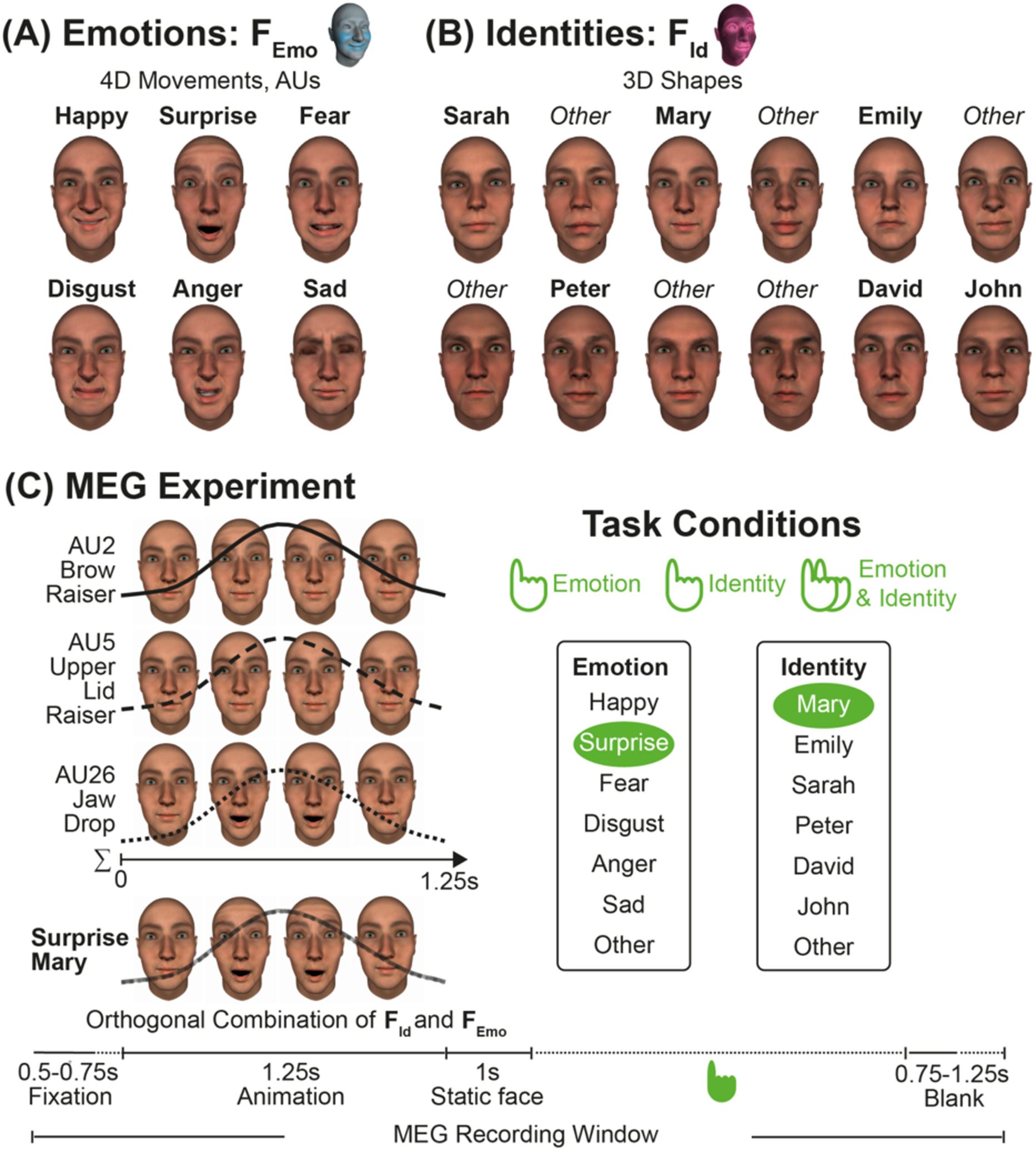
Experimental design. **(A) Emotions.** We generated one facial expression model for each of six classic emotions–happy, surprise, fear, disgust, anger, and sad–using existing F_Emo_ data ^20,29^ (see *Methods: Stimuli*). **(B) Identities.** We generated twelve random face identities using F_Id_ data (see *Methods: Stimuli*). We trained each participant to recognize a random subset of six identities by first name (“Known”) and to label the other six as “other” (“Unknown”). **(C) MEG experiment.** For each emotion model, we rendered 600 animations (3,600 total). Within each animation, we independently sampled the AUs (0 = absent; 1 = dynamically expressed at maximum intensity; see *Methods: Experiment, Stimuli*) and randomly paired these animations with the 12 identities. We randomly assigned participants to one of three task conditions—Identity, Emotion, or Identity & Emotion categorizations. All participants viewed the same 3,600 animations; only the task instructions and response types differed. In the Identity & Emotion task, participants provided two responses per trial, with response order randomized across trials. Each trial began with a 0.5–0.75s fixation cross, followed by a 1.25s dynamic facial animation, then a 1s static face with the response screen, followed by a 0.75–1.25s inter-trial interval (see *Methods: Experiment, Procedure*). We recorded MEG responses throughout.

Next, we generated a stimulus set of 3,600 facial animations by fully crossing the twelve face identities with the six emotion models. For each animation, we independently sampled the AUs defining one of the six emotions and randomly paired the resulting animation with one of the twelve identities (12 identities x 6 emotions x 50 variants, see *Methods, Experiment, Stimuli*). This factorial crossing ensured that identity and emotion features varied independently across trials, preventing stimulus-level covariation between F_Id_ and F_Emo_.

Twenty-four participants viewed the full set of 3,600 animations and performed one of three categorization tasks: (1) identity, (2) emotion, or (3) both identity and emotion (8 participants per task, see *Methods, Participants* and Figure 2). During the experiment, facial animations were presented in a random order while we recorded MEG and behavioral responses (see *Methods, Experiment, MEG Procedure, MEG Data Acquisition and Pre-processing*).

Crucially, all participants viewed the identical set of facial animations and differed only in task instructions. As identity (F_Id_) and emotion (F_Emo_) were orthogonally manipulated, task-demands could selectively prioritize one feature class while sensory input remained constant. This design therefore isolates task-dependent computations in the transformation of facial features into behavior.

By using Mutual Information (MI)^36^, we quantified, across trials, the relationship between the features defining each identity (F_Id_) and emotion (F_Emo_) and the corresponding MEG amplitude response at each source (see *Methods, Analyses, Task-dependent Dynamic Feature Representations*). MI quantifies how strongly neural activity represents a stimulus feature without assuming linearity or specific decoding models^36^. Specifically, we computed MI(F_Id_; MEG) and MI(F_Emo_; MEG) every 4 ms from 0-1000 ms post-animation onset and applied false discovery rate (FDR) correction across 8,196 sources × 250 time points per participant (*p* < .05). This analysis yielded, for each participant and task, a source-by-time matrix of feature-specific neural information, revealing where and when F_Id_ and F_Emo_ are represented across the brain.

As the stimulus features are explicitly parametrized by the generative model, these feature-resolved neural representations can be used to test structured dependencies in cortical processing. Specifically, we examine whether cognitive task demands determine (i) which features are gated, resulting in maintenance or attenuation at early occipital stages, (ii) how maintained features are directionally routed through ventral and lateral pathways, and (iii) whether convergent inputs support conditional nonlinear feature integration in temporal cortex.

### Stage 1: Task-Dependent Feature Gating in OCC

To test the first dependency in our framework (Figure 1; Stage 1 feature gating), we asked whether OCC initially represents both feature classes, and whether task demands determine which feature is maintained longer and which attenuates earlier. If so, this early maintenance-attenuation dynamic (i.e. feature gating) would constrain the availability of information for subsequent routing and integration. We therefore compared OCC feature representations across the three task conditions while holding visual input constant.

To do so, for each participant, and separately for F_Id_ and F_Emo_, we extracted the maximum MI across sources within OCC (including Lingual Gyrus (LG), Pericalcarine (PCAL), Cuneus (CUN), Lateral Occipital Cortex (LOC), see *Methods, Analyses, Stage 1: Occipital Representational Persistence*). These per-timepoint maxima were then averaged across participants within each task group to yield group-level time courses (Figure 3), capturing the temporal evolution of feature-specific information and revealing how task demands determine whether feature information is maintained or attenuated over time.

**Figure 3.**
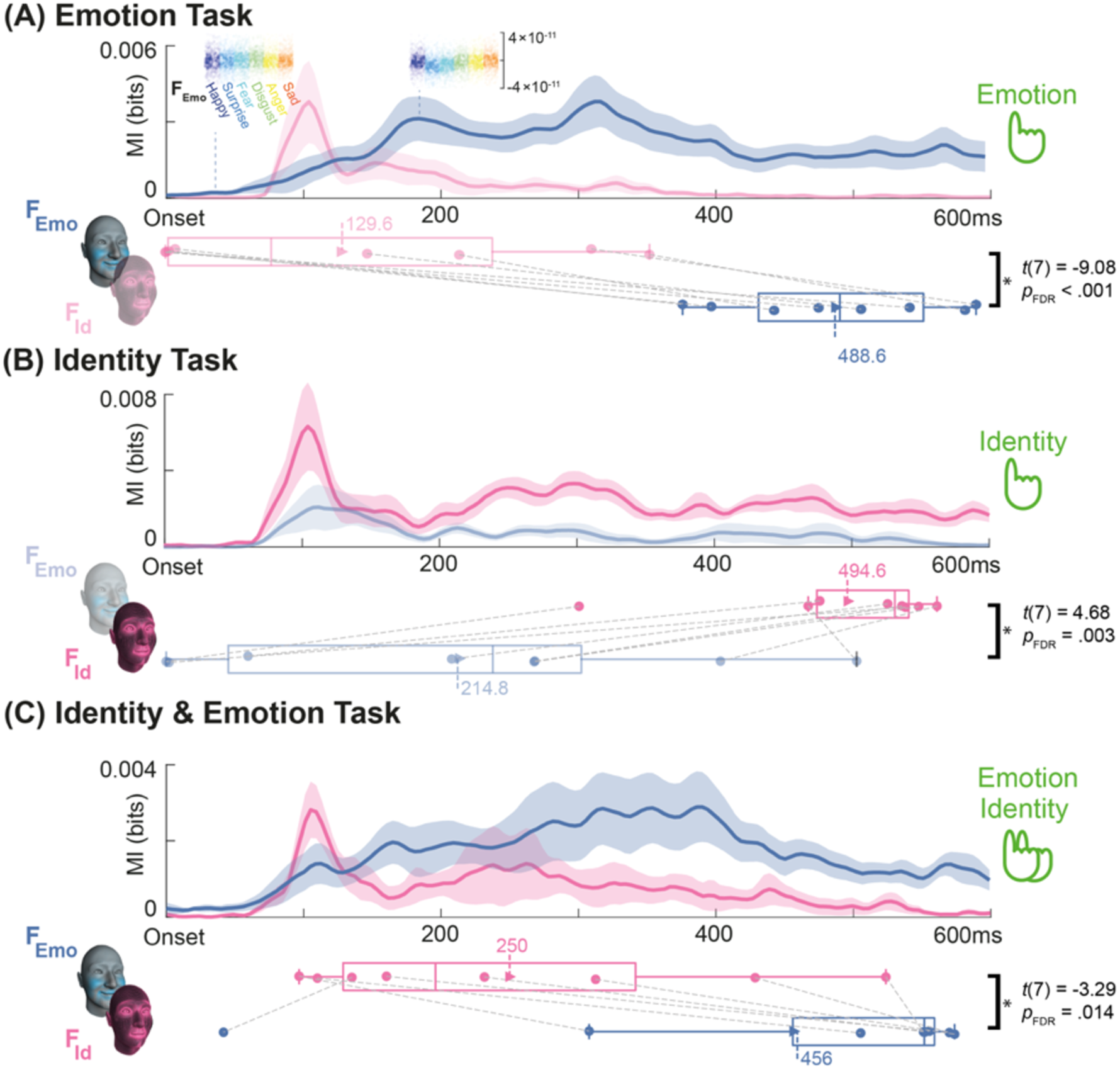
Task-dependent feature gating in OCC revealed by differential maintenance and attenuation of F_Id_ and F_Emo_. We traced the dynamic representations of F_Id_ and F_Emo_ using MI(MEG; F_Id_ or F_Emo_) in OCC. **Panel conventions.** In each panel, the blue curve shows F_Emo_ representations; the magenta curve shows F_Id_ representations (shading = SEM across participants). Curves show the cross-participant average of mean MI (per time point) across OCC sources. Boxplots summarize the distribution across participants of the mean last significant time (averaged across OCC sources) for each feature (F_Id_: magenta; F_Emo_: blue); filled dots indicate individual participants and dashed lines connect paired F_Id_ and F_Emo_ values within participant. **(A) Emotion task:** F_Emo_ is maintained longer than F_Id_ in OCC. **(B) Identity task:** the pattern reverses, with F_Id_ maintained longer than F_Emo_. **(C) Identity & Emotion task:** when both features are task-relevant, both remain significant, indicating passage through the gate, although with asymmetric maintenance durations (F_Emo_ > F_Id_). **Rainbow Insets.** In Panel A, we illustrate how MI quantifies the separability of F_Emo_. Single-trial MEG source amplitudes of a representative participant (Participant 6) are color-coded by the ground truth features. On an OCC source, F_Emo_ MI is low (overlapping responses) at 36 ms and high at 200 ms (responses discriminate F_Emo_).

Importantly, these MI curves do not reflect conventional evoked averages but quantify how well single-trial MEG amplitudes discriminate stimulus features (F_Id_ and F_Emo_) at each time point. Thus, differences between curves directly reflect differences in feature-specific neural information rather than amplitude differences per se. The rainbow insets in Figure 3A illustrate this principle for one participant (Participant 6): an OCC source at an early latency (36 ms) shows overlapping responses across F_Emo_ levels (low MI), whereas later time points (184 ms) show clearer separation across F_Emo_ levels (high MI), demonstrating feature-specific representation (here, F_Emo_).

To quantify the duration of feature maintenance in OCC, we computed, for each OCC source, the latest time point at which MI exceeded the significance threshold (“last significant time point”) and averaged this value across OCC sources for each participant, task and feature (see *Methods, Analyses, Task-dependent Dynamic Feature Representations*). This metric indexes how long feature information remains significantly represented in OCC before attenuating. Figure 3 summarizes the results. A 2 (Feature: F_Id_ vs. F_Emo_) × 3 (Task) repeated-measures ANOVA on this occipital maintenance metric revealed a significant main effect of Feature (*F*(1,7) = 12.18, *p* = .010) and a strong Feature × Task interaction (*F*(2,14) = 32.25, *p* < .001), indicating that occipital feature maintenance depends jointly on feature type and task relevance. Post-hoc paired comparisons within each task (FDR-corrected across the three tests; see boxplots in Figure 3) confirmed task-dependent reversals in which feature is maintained longer than the other.

In the Emotion task, F_Emo_ was maintained longer than F_Id_ (488.6 ms vs. 129.6 ms), indicating relative attenuation of identity information under this task. This effect also replicated at the individual level: using paired *t*-tests across OCC sources’ last significant time points (see Supplementary Table S1), all 8 participants showed longer maintenance for F_Emo_ than F_Id_ (*p* < .05, FDR-corrected across participants; Bayesian Population Prevalence = 1 [0.70, 1.00], a Maximum A Posteriori (MAP) estimate of [95% Highest Posterior Density Interval (HPDI)^42^, see Supplemental Table S1).

In the Identity task, the pattern reversed: F_Id_ was maintained longer than F_Emo_ (494.6 ms vs. 214.8 ms), indicating relative attenuation of emotion information. This reversal was likewise robust at the individual level, with 7/8 participants showing the same effect (BPP = 0.87 [0.54, 0.99], MAP [HPDI], see Supplemental Table S1).

In the Identity & Emotion task, both features remained significant, indicating passage through the gate, although F_Emo_ was maintained longer than F_Id_ on average (456.4 ms vs. 249.5 ms). Notably, the mean last-significant time for F_Id_ (249.5 ms) lies beyond the typical early occipital response peak (i.e., 129.6 ms in the Emotion task), indicating that identity information remains maintained beyond initial representation, although relatively more attenuated than emotion information.

Together, these results show that OCC does not encode both feature classes uniformly: task relevance determines which feature is maintained and which attenuates over time, establishing a gating operation that constrains downstream routing and integration.

### Stage 2: Task-Dependent feature routing through ventral and lateral pathways

Stage 1 established an early occipital gate that determines which feature stream remains available for downstream processing. Stage 2 tests the next information-processing dependency in our framework (Figure 1; Stage 2 routing): once a task-relevant feature survives this OCC gate, does it become selectively represented in higher-level pathways, and does this representation reveal systematic pathway biases consistent with ventral and lateral routing architectures?

### 2.1 Task-relevant features are represented in higher-level regions

We first examined where feature information emerges in higher-level face-processing pathways (Table 1)^23,29,31^, separately for each task (Figure 4, see *Methods, Analyses, Feature Representations in Higher-level Regions*). Figure 4 shows onset maps indicating the first significant F_Emo_ and F_Id_ representations in the cross-participant average. These maps provide a direct test of routing by revealing whether feature information that survives the occipital gate is subsequently represented in higher-level cortex.

**Figure 4.**
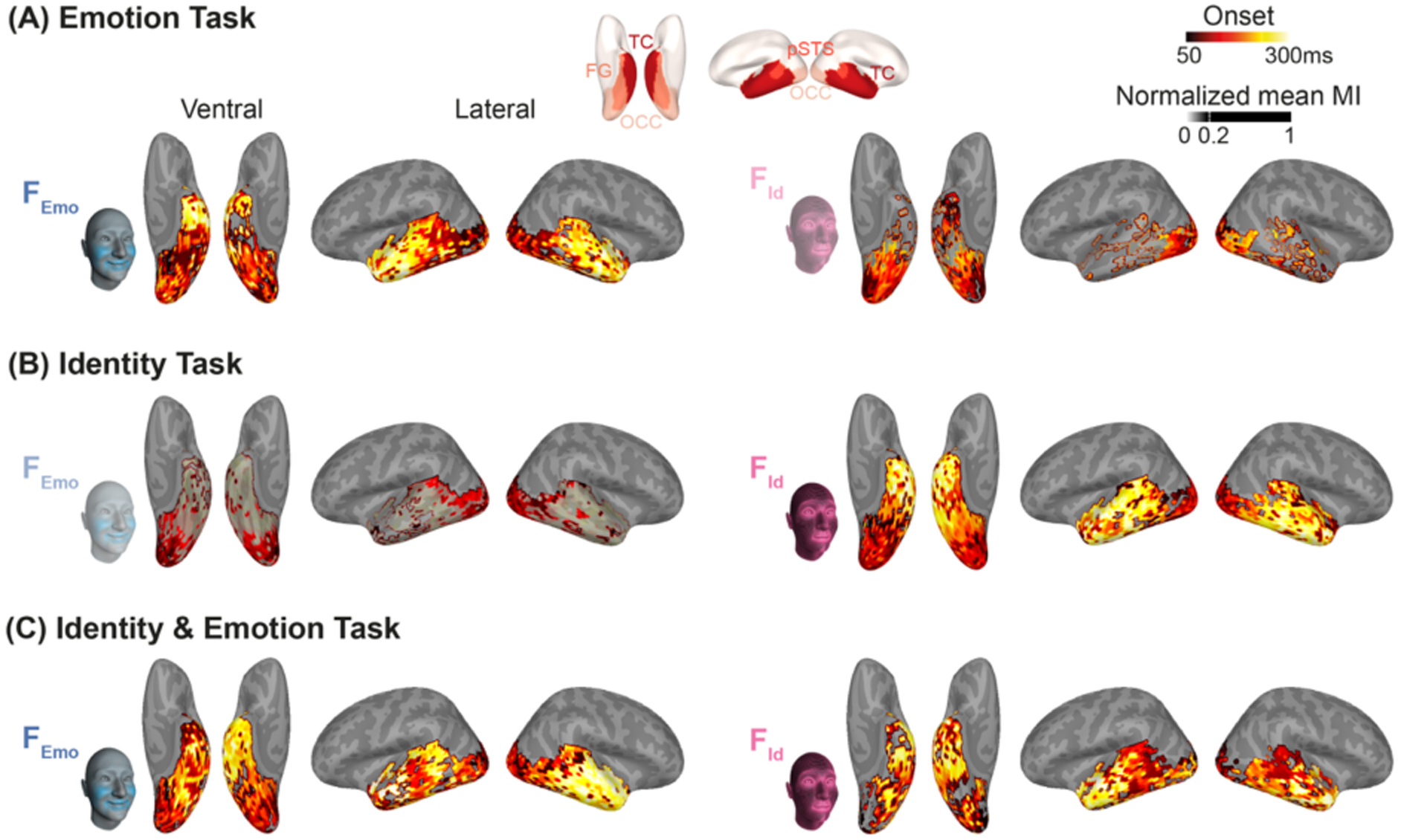
Task-dependent feature onsets into higher-level ventral and lateral pathways. **Panel conventions.** In each panel, surface insets show onset maps for F_Emo_ and F_Id_ (first significant MI-based representations of F_Id_ and F_Emo_ in the cross-participant averages), colour-coded from dark red to yellow (50–300 ms). Transparency is scaled by normalized regional mean MI. Mean MI was first averaged across sources within each region and normalized by the maximum regional value. Values greater than 0.2 were rendered fully opaque, whereas values below 0.2 were mapped directly to transparency. These onset maps indicate when and when feature information becomes represented, not the direction of information flow. **(A) Emotion task.** Task-relevant F_Emo_ is strongly expressed in higher-level pathways and shows OCC-to-ventral and lateral pathway progression, whereas task-irrelevant F_Id_ remains primarily confined to OCC. **(B) Identity task.** The pattern reverses—task-relevant F_Id_ is strongly expressed and extends beyond OCC, whereas task-irrelevant F_Emo_ remains confined to OCC. **(C) Identity & Emotion task**. When both features are task-relevant, both F_Id_ and F_Emo_ show the OCC-to-ventral and lateral pathway progression, enabling subsequent integration.

**Table 1.** Brain regions within the three pathways.

| Pathways | Regions |
| --- | --- |
| Ventral pathway | Fusiform Gyrus (FG)<br>Inferior Temporal Gyrus (ITG) |
| Lateral (third, “social”) pathway | Posterior Superior Temporal Sulcus (pSTS)<br>Middle Temporal Gyrus (MTG)<br>Superior Temporal Gyrus (STG) |

The red (early) to yellow (late) onset maps reveal a clear routing signature: task-relevant feature information progresses from OCC into higher-level cortex, whereas task-irrelevant feature information remains largely confined to OCC. In the Emotion task (Figure 4A), F_Emo_ shows OCC-to-TC progression, whereas F_Id_ remains restricted to the early OCC stage. In the Identity task (Figure 4B), the pattern reverses: F_Id_ progresses beyond OCC, whereas F_Emo_ remains confined. In the Identity and Emotion task (Figure 4C), both F_Id_ and F_Emo_ show OCC-to-TC progression. Together, these onset patterns indicate that feature information is selectively routed into higher-level temporal pathways according to task demands.

### 2.2 Task-relevant features are preferentially represented in the lateral vs ventral nodes

To test whether feature routing is biased toward specific higher-level nodes, we focused on two face-selective targets that occupy complementary positions in the cortical hierarchy: posterior STS (pSTS; lateral pathway) and posterior fusiform gyrus (pFG/Fusiform Face Area (FFA); ventral pathway). These nodes are classically associated with distinct domains–dynamic social signals in pSTS and identity-related structure in ventral fusiform cortex. We therefore asked whether task-relevant emotion features (F_Emo_) preferentially engage pSTS, whereas task-relevant identity (F_Id_) preferentially engage pFG. This provides a direct test of pathway-biased routing at the level of intermediate cortical nodes.

Figure 5 illustrates this node-selective routing (see *Methods, Analyses, Selective Routing of Task-relevant Features*). For task-relevant F_Emo_ (Emotion, and Identity & Emotion tasks; Figure 5A), MI time courses show consistently stronger representations in pSTS than pFG, and participant-level paired comparisons confirm this lateral bias (pSTS > pFG; paired t-test on the max-over-time, mean-over-sources feature representation across participants: *t*(15) = 3.83, *p* = .0016). At the individual level, 15/16 participants replicated this effect (BPP = 0.93 [0.73, 1.00], MAP [HPDI], see Supplemental Table S2).).

**Figure 5.**
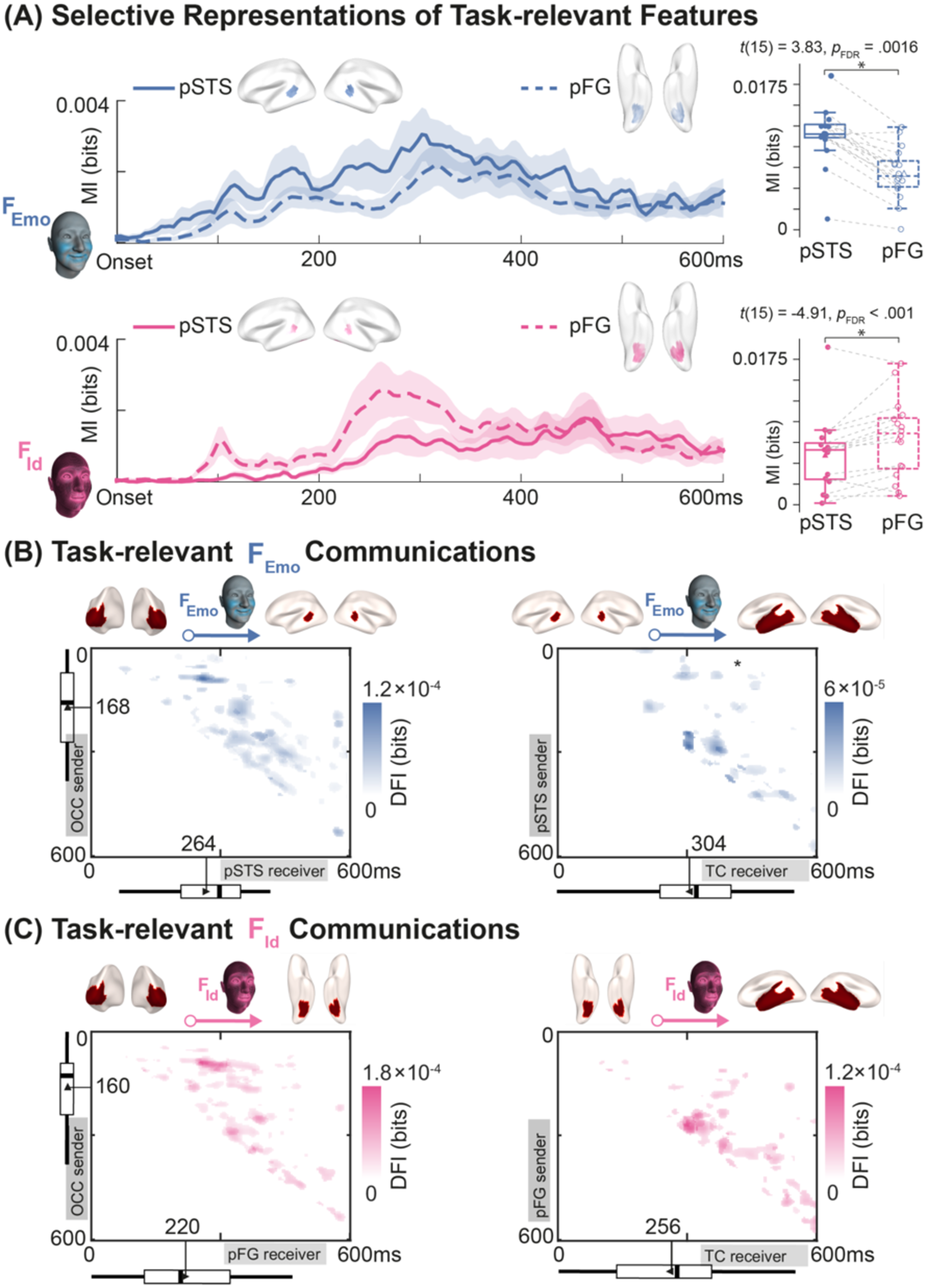
Feature routing and communication in ventral and lateral pathways. **(A) Selective representations of task-relevant features.** Curves show source-mean representations of F_Emo_ (blue) and F_Id_ (pink) in pSTS (solid) and pFG (dashed), from stimulus onset to 600 ms, averaged across participants (F_Emo_: Identity and Identity & Emotion tasks; F_Id_: Emotion and Identity & Emotion tasks). Boxplots compare, across participants, the max-over-time, mean-over-sources feature representation; filled dots indicate individual participants, and dashed lines connect paired values within participant. The boxplots confirm that F_Emo_ is more strongly represented in pSTS than in pFG, whereas F_Id_ is more strongly represented in pFG than in pSTS. **(B) Task-relevant F_Emo_ communications.** For each participant, sender and receiver sources maximally representing F_Emo_ were identified for OCC→pSTS and pSTS→TC pathways. DFI was then computed every 4 ms (0 to 600 ms) across sender (y-axis) and receiver (x-axis) time points, capturing communication delays. Matrix plots show participant-mean significant DFI feature-specific communication (FWER-corrected across 150 × 150 time points, maximum delay 400 ms, *p* < .05). Earliest significant communication shows OCC→pSTS (168->264 ms), followed by pSTS→TC, (→304 ms). **(C) Task-relevant F_Id_ Communications.** Same analysis for OCC→pFG and pFG→TC (Emotion and Identity & Emotion tasks). Earliest significant communication shows OCC→pFG (160→220 ms) followed by pFG→TC (→256 ms). Together, these results demonstrate sequential, feature-specific communication along ventral (F_Id_) and lateral (F_Emo_) pathways. Differences in timing reflect the temporal unfolding of dynamic AU-based features.

By contrast, for task-relevant F_Id_ information (Identity, and Identity & Emotion tasks, Figure 5A), the pattern reverses: F_Id_ representation is stronger in pFG than pSTS, yielding a robust ventral bias (pSTS < pFG; *t*(15) = −4.91, *p* < .001). At the individual level, 13/16 participants replicated this effect (BPP = 0.80 [0.57, 0.94], MAP [HPDI], see Supplemental Table S3).

Together, these results reveal pathway-biased, node-selective routing of facial features: dynamic emotion information is preferentially represented in the lateral pSTS node, whereas identity information is preferentially represented in the ventral pFG node. This establishes that routing is not only expressed at the level of large-scale pathways (Figure 4) but is instantiated at specific intermediate nodes. Importantly, this dissociation reflects feature-biased routing rather than strict anatomical segregation: both nodes contain both feature classes, but task-relevant emotion is preferentially represented in pSTS whereas task-relevant identity is preferentially represented in pFG.

Notably, extending the same comparison to the broader TC revealed no reliable dissociation between ventral ITG and lateral MTG/STG (see Supplemental Table 4), indicating that routing biases are most pronounced at these intermediate pFG–pSTS nodes rather than uniformly across downstream cortex.

### 2.3 Functional communication from OCC to TC via selective nodes

The routing patterns observed above suggest two candidate feature-specific pathways: an emotion pathway extending from OCC to pSTS and onward to TC, and an identity pathway extending from OCC to pFG and onward to TC. To test whether these routes reflect feature-specific communication rather than generic signal coupling, we quantified Directed Feature Information (DFI)^37^, a measure that isolates directed, stimulus-feature specific information transfer (F_Id_ or F_Emo_) between regions, beyond general statistical dependence in MEG signals (see *Methods, Analyses, Task-relevant Feature Communications*).

Using data from the Emotion and Identity & Emotion tasks, for each participant, we identified the sources within sender and receiver regions that maximally represented the relevant F_Emo_ information. We then analyzed feature-specific communication from the OCC to pSTS, and from pSTS to TC, computing DFI_Emo_(sender→receiver) over receiver times from 0 to 600 ms and communication delays spanning 0–400 ms, with FWER correction across 150 receiver time points × 100 delays (*p* < .05). The analogous identity communications DFI_Id_(OCC →pFG) and DFI_Id_(pFG →TC) were computed using the data from Identity, and Identity & Emotion tasks.

Consistent with sequential feature-specific communication along the proposed pathways, the DFI maps (Figure 5B) reveal a temporally ordered progression from OCC to the selective node and subsequently to TC. In Figures 5A-B, F_Emo_ is communicated from OCC at 168 ms, received by pSTS at 264 ms, and subsequently received by TC at 304 ms. In Figure 5C, F_Id_ is communicated from OCC at 160 ms, received by pFG at 220 ms, and subsequently received by TC at 256 ms.

These results demonstrate that the node-selective routing observed above is implemented through feature-specific communication along two cortical pathways: an emotion pathway linking OCC, pSTS, and TC, and an identity pathway linking OCC, pFG and TC. Critically, this establishes that routing reflects the directed communication of stimulus-defined information, rather than co-activation or undirected coupling between regions. Together with the regional representation and node selectivity results above, these findings show that task-relevant facial features are not only represented in pathway-specific nodes but are actively and sequentially communicated through them toward downstream temporal systems. These routed feature streams constitute the inputs to the nonlinear feature-integration computations tested in Stage 3.

### Stage 3: Integration of F_Id_ and F_Emo_ into Person-Relevant Meaning

Having established in Stage 1 that task demands determine which feature information is maintained in OCC, and in Stage 2 that maintained features are directionally routed through feature-selective ventral and lateral nodes, we next test the final dependency in our framework: whether TC integrates identity (F_Id_) and emotion (F_Emo_) information into person-relevant meaning.

Our model predicts that such integration should occur only when both feature classes converge in TC following earlier routing (Figure 6A). This prediction implies two constraints. First, identity information must be semantically grounded to support person-level representations. Second, when identity and emotion features converge locally in TC, their joint representation should exhibit nonlinear (synergistic) integration consistent with feature-level computatins, rather than independent co-representation. Demonstrating such conditional integration would establish that person-level meaning arises from structured feature convergence under these constraints, rather than from independent processing streams.

**Figure 6.**
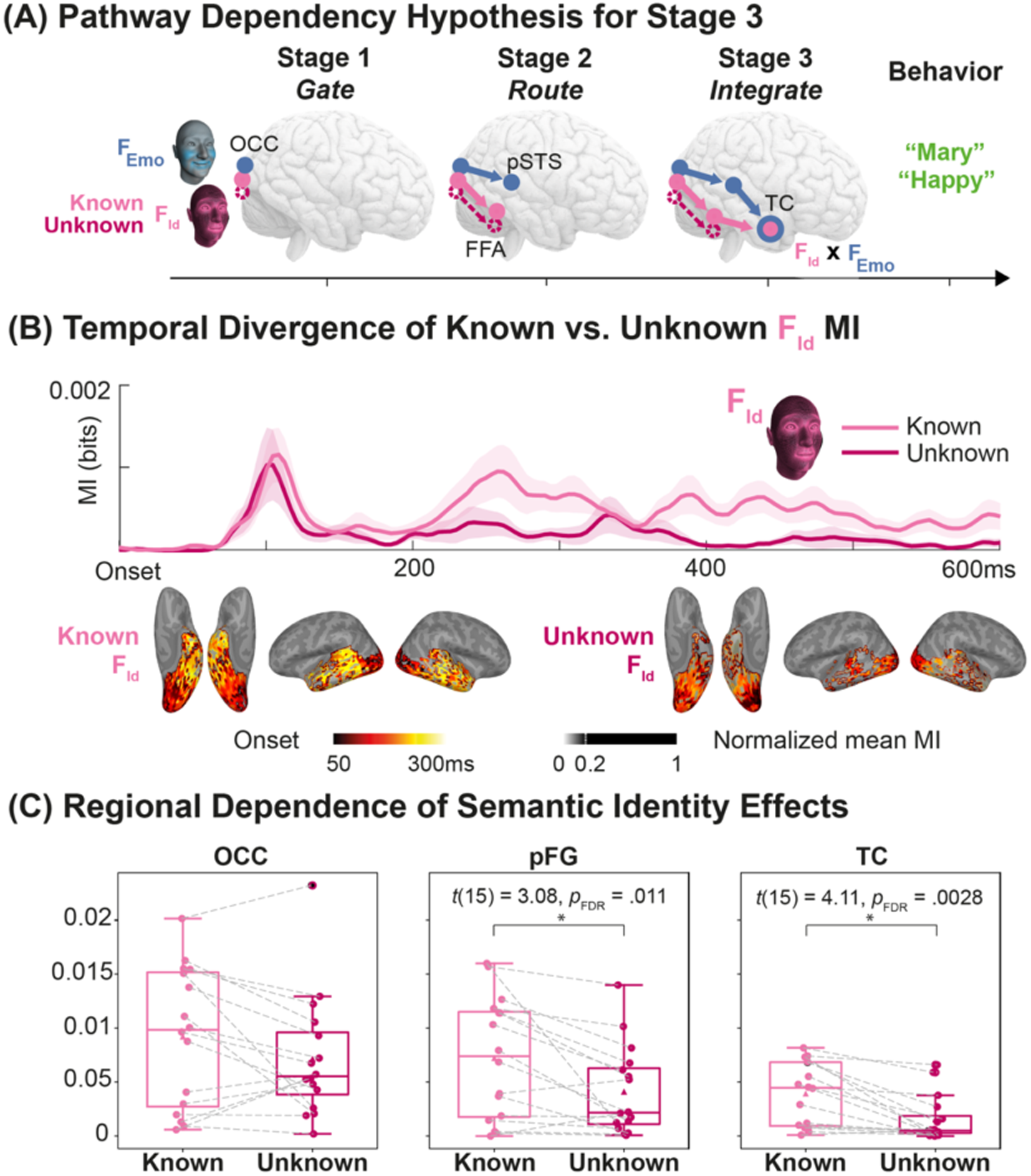
Identity reaches temporal cortex only when semantically known. **(A) Pathway dependency hypothesis for stage 3 integration.** Following Stage 2 routing identity (F_Id_) and emotion (F_Emo_) features converge in TC. We test whether F_Id_ contributes to this convergence only when semantically grounded (Known, light pink) versus not (Unknown F_Id_, dark pink) in the Identity and Identity & Emotion tasks. **(B) Temporal Divergence of Task-relevant F_Id_ MI**, Curves show source-mean feature MI representation time courses for Known (light pink) and Unknown (dark pink) F_Id_ from stimulus onset to 600 ms (Identity & Emotion task). Known and Unknown FId overlap in their early OCC representations but diverge at later stages. Surface onset maps illustrate the spatial progression of F_Id_ representations. **(C) Regional dependence of semantic identity effects.** Boxplots show group-level max-over-time, mean-over-sources feature MI for Known and Unknown F_Id_ in OCC, pFG, and TC. No reliable difference is observed in OCC, whereas Known F_Id_ representation is significantly stronger than Unknown F_Id_ in pFG and TC. This pattern demonstrates that identity information reaches TC in a semantically usable form only when the identity is known.

We therefore first tested whether identity propagation to TC depends on semantic knowledge of the person, and then whether convergent identity and emotion features give rise to integrated neural representations. This directly tests the final prediction of our model: that person-level meaning emerges only when maintained features are routed and subsequently integrated, rather than as an automatic consequence of stimulus exposure or feature co-occurence.

Stage 3 focuses specifically on the dual-task condition, in which both feature classes are behaviorally relevant and therefore available for testing feature convergence and integration. We treat TC as a convergence territory rather than further subdividing it into ventral and lateral substreams because the strongest feature-selectivity dissociation was observed earlier at the intermediate pFG-pSTS routing nodes.

### 3.1 Identity reaches temporal cortex only when known

To determine whether identity information reaches TC irrespective of semantic status, we compared the propagation of Known and Unknown identity representations along the ventral pathway and into TC in the Identity and Identity & Emotion tasks (see *Methods, Analyses, Temporal Divergence of Known vs. Unknown identities*). Here, Known identities were defined as faces associated with trained names, whereas Unknown identities were physically matched faces that lacked semantic labels for a given participant.

Figure 6B and C show the participant-mean MI (MEG; Known F_Id_) and MI (MEG; Unknown F_Id_), averaged across OCC, pFG and TC sources. A 2 (Known vs Unknown F_Id_) × 3 (Region) repeated-measures ANOVA on the max-over-time, mean-over-sources metric revealed a significant main effect of Feature (*F*(1,15) = 8.42, *p* = .011) and Region (*F*(2,30) = 18.30, *p* < .001), with no Feature × Region interaction (*F*(2,30) = 1.25, *p* = .30). Planned paired comparisons within each region showed that the known–unknown difference was significant in pFG (*t*(15) = 3.08, *p*_FDR_ = .011) and TC (*t*(15) = 4.11, *p*_FDR_ = .0028). At the individual level, the effect was less stable in pFG, replicating in 10/16 participants (BPP = 0.61 [0.36 0.81], MAP [HPDI], see Supplemental Table S5), but highly consistent in TC, replicating in 14/16 participants (BPP = 0.89 [0.65, 0.98], MAP [HPDI], see Supplemental Table S6). Together, these results indicate that identity information reaching TC depends on semantic knowledge of the person.

This distinction imposes an important constraint on the computational logic of Stage 3: identity information that reaches TC is not purely visual but selectively associated with semantic knowledge. If TC simply inherited identity-related visual information from the ventral pathway without further constraint, Known and Unknown identities would be expected to show comparable downstream propagation once represented in pFG. Instead, Known identity was represented more strongly and more consistently than Unknown identity in TC, indicating that semantic knowledge gates or stabilizes the propagation of ventrally routed identity information to downstream convergence sites. In other words, Stage 2 routing is necessary but not sufficient for Stage 3 integration: identity information must also be semantically interpretable to support robust person-level representation. Thus, identities linked to learned person knowledge are preferentially propagated along the ventral pathway, making them more available for convergence with routed emotion information in TC.

This establishes a representational bottleneck at the entry to Stage 3: only semantically grounded F_Id_ can participate in subsequent feature integration with F_Emo_.

### 3.2 Integration of Known F_Id_ and F_Emo_ in temporal cortex

We examined this process in the Identity & Emotion task, when both F_Id_ and F_Emo_ were task-relevant, identities were semantically known, and emotions were correctly categorized (Figure 6A). Analyses focused on TC where ventral and lateral pathway features converge. Integration refers to neural representations in which F_Id_ and F_Emo_ jointly determine local MEG activity non-additively, rather than varying independently.

Because integration can only occur where both features are locally represented, we restricted the analysis to regions and timepoints showing significant MI(F_Id_; MEG) and MI(F_Emo_; MEG)–that is, where ventral and lateral inputs had converged. Under this condition, Synergy quantifies complementary, non-redundant contributions of F_Id_ and F_Emo_ to the neural signal, consistent with nonlinear integration rather than additive co-representation.

For each participant and each of the 792 sources within TC, we computed Synergy(F_Id_; F_Emo_; MEG) every 4 ms from 0-600 ms post-stimulus onset. Higher synergy values indicate encoding of the joint (F_Id_ x F_Emo_) representation beyond either feature alone (see *Methods, Analyses, Feature Integration*).

To illustrate this computation, Figures 7B and C show data from a representative participant. The mean MI time courses for F_Id_ and F_Emo_, averaged across TC sources, peaked 68 ms apart, consistent with the temporal unfolding of facial movements in the stimuli. The source exhibiting maximal Synergy (Figure 7C) peaked at 316 ms, after both features were locally available, confirming that integration occurred once F_Id_ and F_Emo_ were locally represented.

**Figure 7.**
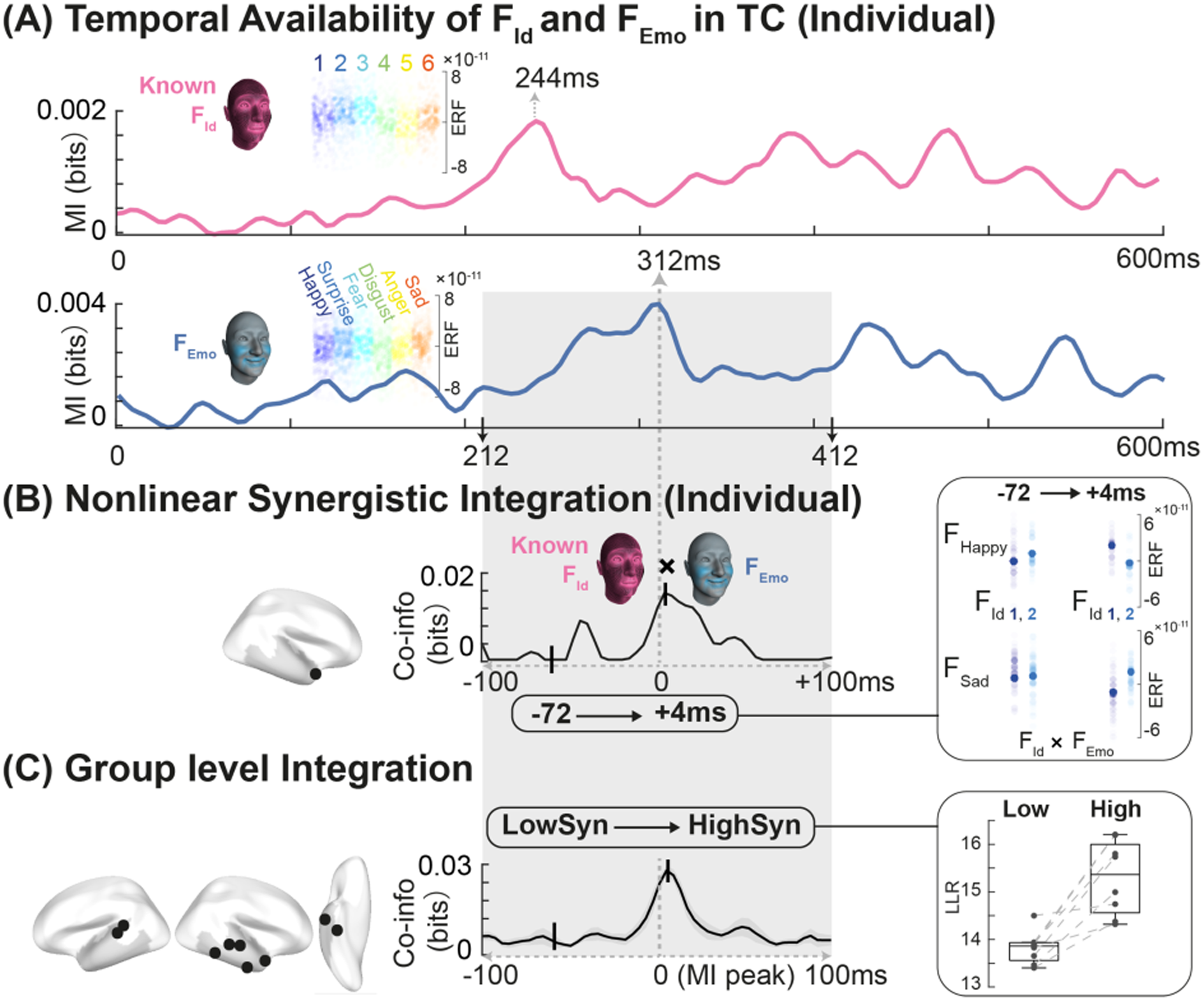
Nonlinear Integration of F_Id_ and F_Emo_ in TC. **(A) Temporal availability of F_Id_ and F_Emo_ in TC (individual participant).** Mean MI timecourses show F_Id_ (pink) and F_Emo_ (blue) representations in TC during the Identity & Emotion task. FId and FEmo peak at different latencies (244 ms and 312 ms), establishing a temporal offset that constrains their joint availability. The grey region marks the analysis time window. Insets illustrate single trial separability at peak MI. **(B) Nonlinear synergistic integration at the source level (Individual).** Synergy(F_Id_; F_Emo_; MEG) was computed for each TC source (4 ms resolution, 0–600 ms). The time course corresponds to the source with maximum synergy (black dot on cortex), realigned to the later of the two MI peaks (0 ms). Synergy rises immediately after this point, indicating that F_Id_ and F_Emo_ integration occurs only once both features are locally represented. Insets illustrate this integration: at −72 ms (low Synergy), F_Id_ separation (1 vs. 2) is invariant across F_Emo_ (happy vs. sad); at +4 ms (high Synergy), F_Id_ separation depends on F_Emo,_ demonstrating joint representation. **(C) Group level integration.** The same alignment was applied across participants (N = 8; F_Emo_ later peak in 7/8 participants). The mean synergy (± SEM) increases sharply around 0 ms, confirming that TC systematically follow ventral (F_Id_) and lateral (F_Emo_) feature convergence in TC. Cortical renderings display each participant’s TC source with maximum synergy (black dots). Boxplots show the model-based validation of non-linear integration. Log-likelihood (LLR) distributions compare models with and without an interaction term (F_Id_ x F_Emo_). Higher LLR at peak (vs. low) synergy confirms that TC activity reflects genuine nonlinear feature interaction rather than additive co-representation. Small dots mark individual data points; grey lines connect within-participant LLRs across low and high synergy time point.

Insets in Figure 7C further illustrate this logic: at −72 ms (low Synergy), separation between F_Id_ (1 vs. 2) was similar across F_Emo_ (happy vs. sad); at +4 ms (high Synergy), separation of the same F_Id_ varied with F_Emo_, demonstrating feature-dependent joint coding.

At the group level (Figure 7C), we related each participant’s synergy time course to the later of the two MI peaks (F_Emo_ in 7/8 participants). Across participants, Synergy peaked immediately after this point, demonstrating that integration follows–and therefore depends on–feature convergence in TC.

To confirm that Synergy reflects genuine feature interaction, we compared a baseline model (main effects only) with a full model including the interaction terms (F_Id_ × F_Emo_) (see *Methods, Analyses, Feature Integration*). The interaction contribution, quantified by a log-likelihood ratio (LLR), increased significantly at maximum vs. minimum Synergy (mean LLR difference = 10.68 ± 5.99, *t*(7) = -5.04, *p* = .0015), confirming that the interaction term explains additional variance specifically when Synergy is high.

Extending this analysis across TC (including for each participant their ITG, MTG, and STG sources, see Methods, Analyses, Feature Integration) yielded the same effect *t*(7) = -5.15, *p* = .001, Figure 7D, boxplot). This effect replicated across all participants (paired *t*-tests across each participant’s sources, *t*(791), *p* < .05, BPP = 1 [0.70 1.00], MAP [95% HPDI]).

Together, these results demonstrate that TC encodes a higher-order representation arising from nonlinear synergistic integration of F_Id_ and F_Emo_. Because this code emerges only when identities are semantically known and emotions correctly categorized, it reflects semantic-level integration rather than perceptual co-activation. Thus, TC combines routed feature inputs into a unified, person-relevant representation (e.g. “happy Mary”), marking the transition from perceptual processing to person-level meaning.

Taken together with Stages 1 and 2, these results establish a strict dependency structure: task demands gate feature availability (Stage 1), maintained features are then selectively routed (Stage 2), and only when routed features converge and satisfy semantic constraints are they nonlinearly integrated into person-relevant meaning (Stage 3). Flexible person perception therefore arises from sequential, task-dependent computational constraints on feature availability, communication, and integration–not from passive feature accumulation.

## Discussion

We reveal a structured information processing architecture of cortical computations that transforms facial features into person-relevant meaning. Task demands first gate which facial features are maintained in occipital cortex, then selectively route these features through ventral and lateral pathways via feature-specific communication, and only then permit their nonlinear integration in temporal cortex when semantic identity information is available.

Using generative 4D face modelling, participant-specific millisecond-resolved MEG, and information-theoretic and model-based analyses, we traced the sequential feature-level computations operating at the algorithmic level ^38^ that gate, communicate and conditionally integrate identity and emotion features during face tasks. From these results, three computational principles emerge.

First, task relevance actively gates feature maintenance: both identity (F_Id_) and emotion (F_Emo_) are initially represented in OCC, but only task-relevant features are sustained beyond this early stage, whereas identical but task-irrelevant features rapidly attenuate. Second, maintained features show pathway-biased routing: identity information is preferentially communicated through the ventral pathway, whereas emotion information is communicated through the lateral pathway. Third, integration occurs only when routed inputs converge under appropriate semantic conditions. In temporal regions, identity and emotion features combine into a nonlinear (synergistic) code that carries more information than either feature alone. Critically, this convergence emerged only for semantically known identities, indicating that semantic knowledge acts as a constraint on higher-order feature integration.

### Extending models of face perception: from pathway specialization to feature routing

Classical models of face perception–from Bruce and Young’s functional framework^43^ to Haxby’s distributed system^44^–identified specialization for identity and dynamic expression. Neuroimaging and neurophysiology subsequently established anatomico-functional distinctions between the ventral and lateral pathways supporting these computations^24–27,30^. However, these models left unresolved how these pathways are dynamically coordinated during task-dependent processing at the level of feature-based computations operating on facial features.

Our results extend this framework by introducing three computational principles linking perceptual features to meaning. First, task-dependent gating in OCC determines which feature information remains available for subsequent cortical processing, establishing an early computational bottleneck. This finding indicates that face perception operates as an active, goal-dependent computation rather than a passive feedforward cascade. Second, selective routing organizes maintained features across cortical pathways. Static structural features underlying identity are communicated through ventral regions optimized for 3D shape information, whereas dynamic motion features supporting emotion judgments are communicated through the lateral pathway specialized for socially relevant facial dynamics. Third, conditional integration in TC identifies where segregated pathways converge. Only when identity representations are semantically grounded do ventral identity features combine with lateral emotion features to form integrated person-level representations. This conditional convergence provides a feature-level account of how distinct perceptual dimensions extracted from the same face–who someone is and how they feel–become unified within a coherent representation of a person. Importantly, our results support feature-biased routing rather than strict anatomical pathway exclusivity.

### Semantic knowledge as a gate for feature integration

A central finding is that identity-emotion integration depends on semantic familiarity. Only identities associated with semantic labels (Known) are propagated beyond FG to participate in integration within TC. Semantic knowledge therefore acts as a selective gate enabling convergence between perceptual pathways. Without a nameable representation, ventral identity features remain confined to FG and do not combine with lateral emotion representations. This finding aligns with evidence showing that person knowledge modulates TC activity and supports linking perceptual input to conceptual meaning^12,44^, with implications for social cognition and biases^45^ in trait inference from faces ^1–3,5–9,46^. Our results extend this view by revealing the feature-level and temporally resolved dynamics through which such integration emerges.

At the level of social neuroscience, the synergistic F_Id_ × F_Emo_ representation we observe in TC constitutes a distinct representational level. It binds identity and emotional state into a single, feature-compositional semantic construct. This mechanism offers a possible neural basis for a long-standing problem in social perception: how separable pathways for identity and expression converge to support higher-order judgments such as trustworthiness, dominance, or affiliation.

### Attention and the limits of feature integration

Although task-dependent gating resembles attentional modulation, our design differs from classical spatial or feature-based attention paradigms. Identity and emotion features were spatially and temporally co-localized within the same dynamic stimulus, preventing simple spatial or temporal selection. Instead, task demands determined which features were maintained and routed through distinct cortical pathways. Moreover, the synergistic representations observed in TC reflect the emergence of new representational content rather than simple response gain^47^, indicating integration rather than modulation.

### Top-Down influences on early feature gating

The early gating observed in OCC likely reflects rapid top-down control^48,49^. Prior work shows that feedback from prefrontal and parietal regions can modulate OCC representations within 150 ms, selectively enhancing task-relevant features^33,48,50^. Although we did not directly test this, such rapid recurrent interactions could provide a plausible mechanism by which identical visual input is flexibly routed through ventral or lateral pathways depending on task demands. Within this framework, OCC acts as a dynamic interface between sensory input and goal-directed cortical processing, where early feature representations are selectively stabilized or attenuated before being communicated through higher-level pathways.

### Methodological advances and broader applicability

Beyond these theoretical insights, our study introduces a methodological framework for tracing the transformation of perceptual features into higher-order representations. First, generative 4D face modelling provided precise control over ground truth identity and emotion features in dynamic stimuli, avoiding the confounds inherent in naturalistic images or videos where multiple dimensions covary. Second, information-theoretic analyses allowed us to quantify how these features are represented, communicated and integrated over space and time. Mutual Information captured feature-specific representations^36^, while synergy quantified nonlinear integrative codes that exceed additive feature contributions^51^. Together, these tools provide a general framework for studying how cortical systems transform sensory features into behaviorally relevant representations.

Importantly, this framework extends beyond faces. Similar algorithmic principles may govern object and scene perception (integration of local and global features), language (phoneme-prosody coupling), or multimodal social signals (face–voice integration). In each case, task demands may determine which features are maintained, how they are routed and communicated across cortical pathways, and when they converge to produce emergent meaning^52^.

### Limitations and future directions

Several limitations warrant consideration. First, while MEG provides millisecond resolution of cortical dynamics and is well suited to characterizing feature-level computations at the algorithmic level, spatial precision remains limited. Combining E/MEG with laminar fMRI or electrocorticography, could refine anatomical localization and test circuit-level (laminar) implementation of feature gating, routing, communication and integration. Second, our stimuli represented Western White identities and six canonical expressions. Extending this approach to more diverse identities, cultures and expressions will be important to test the generality of these mechanisms. Third, person perception is inherently multimodal^39,43,44^.

Future studies integrating visual, auditory, and contextual modalities could examine how TC integrates information across modalities to construct person meaning. Finally, the synergy framework raises broader questions about neural compositionality. Do similar mechanisms combine gaze, prosody, or gesture into unified representations of intent and affect? Addressing these questions may reveal general computational principles underlying social cognition.

### Bridging brain and artificial intelligence

Our findings also inform debates in artificial intelligence, particularly the symbol-grounding problem–how abstract meaning becomes anchored in sensory input. The cortical architecture identified here suggests a biological solution: task demands gate feature selection, pathways route and communicate feature-specific information streams, and TC integrates them into nonlinear synergistic representations grounded in perception.

These principles suggest design directions for more flexible and interpretable AI systems^53–55^. Contemporary deep neural networks typically process features indiscriminately and lack interpretable mechanisms for dynamic, task-dependent routing. Incorporating selective gating, pathway-specific processing, and explicit nonlinear compositional integration may therefore enable AI systems that generalize more flexibly while maintaining interpretable feature representations. Critically, this perspective emphasizes that alignment between brain and artificial systems require algorithmic-level equivalence–shared feature representations, communications and computations–not merely similarity in outputs ^53^. Such biologically informed models could improve not only performance but also explainability.

## Conclusion

In summary, we identify a structured sequence of cortical computations that dynamically transforms visual features into social meaning. Task demands gate feature maintenance, cortical pathways route and communicate identity and emotion information, and temporal cortex nonlinearly integrates these features when semantic conditions allow. By tracing the spatiotemporal trajectory of stimulus features through the brain, we reveal the systems-level information processing architecture underlying cognition and constrain the computations that support flexible visual categorization.

### Methods

### Participant

Twenty-four participants (12 female participants, Western White, aged 18-35 years, mean = 23.8, SD = 4.3) took part in the experiment and provided informed consent. All had normal or corrected-to-normal vision and reported no history of disorders affecting social signal processing (e.g., autism spectrum disorder), face perception (e.g., prosopagnosia), learning difficulties, synesthesia, or other psychiatric/neurological disorders. The study was approved by the University of Glasgow College of Medical, Veterinary & Life Sciences Ethics Committee (Application 200230234).

## Experiment

### Stimuli

We used a generative model of human facial movements^35^, comprising a library of 42 individual facial action units (AUs)—i.e., the basic elements of human facial movements as detailed by the Facial Action Coding System (FACS)^41^. Each AU in the generative model was derived from real humans, trained to accurately produce each individual AU on their face, captured using a stereoscopic recording system, and whose renderings were verified by the trained AU producers^35^. Therefore, the generative model produces physiologically valid representations of real human facial movements and comprises no physiologically impossible facial movements. For the present experiment, we defined six emotion models (happy, surprise, fear, disgust, anger, sad). For each emotion, we selected the most prevalent AUs from a corpus of 50 person-specific models ^20,29^, as shown in Table 2.

**Table 2.** Emotion models with their AUs.

| Emotion | AUs |  |  |
| --- | --- | --- | --- |
| Happy | Cheek Raiser (AU 6) | Lips Part (AU 25) | Lip Corner Puller (AU12) |
| Surprise | Outer-Inner Brow Raiser (AU1-2) | Upper Lid Raiser (AU5) | Mouth Stretch (AU27i) |
| Fear | Upper Lid Raiser (AU5) | Lower Lip Depressor (AU16open) | Lip Stretcher (AU20) |
| Disgust | Nose Wrinkler (AU9) | Cheek Raiser (Right) (AU6R) | Lip Stretcher (AU20) |
| Anger | Nose Wrinkler (AU9) | Lower Lip Depressor (AU16open) | Lid Tightener (Right) (AU75) |
| Sad | Brow Lowerer (AU4) | Chin Raiser (AU17) | Eyes Closed (AU43) |

Within each animation, we sampled the three AUs independently: on 45% of trials, an AU’s intensity was set to 0 (absent), and on 55% of trials it was set to 1 (present at maximum). To deliver natural perception, five temporal parameters—onset latency, acceleration, peak latency, deceleration, and offset latency—were fixed at 0.5. Together with the intensity parameter, these six controls yielded a single-peaked trajectory for each visible AU, reaching its maximum at 0.63 s within a 1.25 s animation.

We manipulated identity using a 3D face identity generator ^15^ to create 12 random identities (Western White, 6 female, 6 male, aged 25 years). Face identities were generated using a 3D generative face model (GMF) built from a database of 355 scanned faces. A general linear model decomposed each face into categorical factors (sex, age, ethnicity) and identity-specific residuals. By holding categorical factors constant and randomizing the residual identity components (via PCA), the model synthesized novel random 3D identities.

In total, we rendered 3,600 facial-expression animations (600 per emotion), each paired at random with one of the 12 identities. In the MEG scanner, participants maintained a constant viewing distance of 115 cm, with stimuli subtending 14° (vertical) × 10.5° (horizontal) of visual angle to simulate conversational distance. All participants viewed the same 3,600 animations.

### Training Procedure

Before MEG, participants were trained to recognize both identities and emotions to a high accuracy criterion. Each participant was assigned 6 of the 12 identities and given specific names for these (Emily, Sarah, Mary, Peter, David, John); the remaining six identities had no assigned name and were to be answered as “Other.” Training proceeded in short blocks with immediate feedback. At the start–and again before each block–we showed the six named faces with their labels. On each identity trial, participants viewed a random animation from the stimulus set and then chose a response from six names plus “Other.” The first two blocks contained only the six named faces; from Block 3 onward, 50% of trials presented unnamed faces for which “Other” was the correct response. Emotion training followed the same format but used animations from the full emotion models (all three AUs visible) and required participants to select among the six emotion labels. Participants continued until they achieved an accuracy of >95% in two consecutive blocks (for identity, criterion assessed from Block 3, once “Other” trials were present). All participants completed both training procedures within one hour, reached the criterion, and reported no confusion between labels and animations.

### MEG Procedure

After training, participants completed MEG sessions under a pre-assigned task: Identity, Emotion, or Identity & Emotion. On each trial, participants viewed one of the 3,600 animations (the same stimuli set for all participants, presented in a randomized order) and responded using the same options as in training. In the Identity task, the options were Emily, Sarah, Mary, Peter, David, John, or Other; in the Emotion task, they were Happy, Surprise, Fear, Disgust, Anger, Sad, or Other. In the Identity & Emotion task, participants gave both responses, with the order of Identity vs. Emotion responses randomized per trial.

Each trial started with a jittered fixation period (0.5 s–0.75 s), followed by a 1.25s face animation and then the same static face for 1 s to maintain continuity. The response screen then appeared, followed by an inter-trial interval (ITI) of 0.75-1.25 s (jittered). A central fixation cross was present throughout except during the ITI. Each participant completed 75 blocks of 48 trials across 3–5 sessions over 3–5 days, yielding a total of 3,600 trials per participant, with MEG recorded continuously.

### MEG Data Acquisition and Pre-processing

We measured each participant’s MEG activity with a 306-channel Elekta Neuromag MEG scanner (MEGIN) at a sampling rate of 1,000 Hz and tracked continuous head position with cHPI. We performed the analyses according to recommended guidelines using MNE-python software^56^ and in-house Python/MATLAB code.

We identified and excluded bad channels using a combination of automated detection and visual inspection, and discarded blocks with head motion > 0.6 cm (as measured by cHPI). To suppress environmental noise and compensate for residual motion, we applied Maxwell filtering with signal-space separation (SSS)^57^. We band-pass filtered the data between 1 and 150 Hz (Hamming FIR filter), and notch-filtered the data at 50, 100, and 150 Hz. We rejected muscle bursts and step/jump artefacts with automatic detection procedures. We epoched the data into [0–1000 ms] trial windows around animation onset. Within each session, we concatenated epoched data and decomposed them with ICA; 2–5 components corresponding to ocular and cardiac artefacts were identified and removed.

For analyses, we resampled the output data at 250 Hz, low-pass filtered them at 25 Hz (5th-order Hamming FIR filter). We computed MEG source estimates over time using a minimum-norm estimate (MNE) with noise covariance from empty-room recordings and a participant-specific boundary element model (BEM, computed with FreeSurfer and MNE), evaluated on a 5 mm source grid (regularization parameter *λ* = 1/9). We applied this reconstruction to each session of trials. These computations produced, for each participant, a matrix of single-trial MEG response time series with dimensions 8,196 MEG sources x 250 Hz sampling rate.

## Analyses

### Task-dependent Dynamic Feature Representations

To examine how task relevance shapes the dynamics of Identity (F_Id_) and Emotion (F_Emo_) representation, we computed mutual information (MI) between stimulus features and MEG source responses. For each participant, MI(F_Id_; MEG) and MI(F_Emo_; MEG) were evaluated every 4 ms from 0–1000 ms after animation onset, on each of 8,196 sources.

MI was estimated using the Gaussian Copula Mutual Information (GCMI) method^36^. Statistical significance was assessed with the Benjamini–Hochberg procedure, controlling the false discovery rate (FDR) at *q* < .05 (one-tailed) across 8,196 sources x 250 time points^58^. To obtain empirical p-values for the individual tests, we exploited the fact that the null distribution of GCMI depends only on the dimensionality of the responses and the number of samples per class. This null distribution was generated by shuffling trial labels 1,000,000 times, providing empirical p-values for each source and time point estimate.

### Stage 1: Occipital Representational Maintenance

For each task group, we summarized occipital representational dynamics by (i) taking, at each time point, the mean MI across the 625 sources in OCC (Table 1) and (ii) averaging these values across participants, computed separately for F_Id_ and F_Emo_. We further quantified occipital representational persistence by identifying, for each participant and each source, the last post-stimulus time point at which MI exceeded the significance threshold, computed separately for F_Emo_ and F_Id_. These last significant times were then averaged across OCC sources to obtain a participant-level estimate of persistence for each feature. Group-level differences between F_Emo_ and F_Id_ were assessed within each task using paired *t*-tests across participants, with *p* values FDR-corrected across tasks. At the individual level, F_Emo_ and F_Id_ were further compared using paired *t*-tests across OCC sources within each participant, and these participant-wise *p* values were FDR-corrected across participants within each task. Figure 3 shows the time courses and persistence comparisons, contrasting task-relevant and task-irrelevant features.

### Stage 2: Functional Communications via Selective Nodes

#### Feature Representations in Higher-level Regions

To visualize the spatiotemporal progression of feature representations, we derived onset maps by identifying, for each source in OCC and along the ventral and lateral pathways (Table 1), the first time point at which the participant-mean MI time course exceeded the significance threshold, separately for F_Id_ and F_Emo_. Figure 4 shows the onset maps, contrasting task-relevant and task-irrelevant features.

#### Selective Routing of Task-relevant Features

To compare the distribution of task-relevant feature information across higher-level temporal cortex, we focused on a lateral node (pSTS) and a ventral node (pFG; restricted to selected fusiform subregions). For each feature, we first combined the two task conditions in which that feature was relevant (F_Emo_: Emotion and Identity & Emotion; F_Id_: Identity and Identity & Emotion) and averaged them within participant, yielding one participant-level MI dataset per feature. For each source and feature, we took the maximum MI across 0–600 ms from animation onset. These source-wise values were then averaged within each node to obtain participant-level estimates for pSTS and pFG representations, separately for F_Emo_ and F_Id_. We assessed these data using a 2 (feature: F_Emo_ vs. F_Id_) × 2 (node: pSTS vs. pFG) repeated-measures ANOVA with within-subject factors, followed by planned paired t-tests comparing pSTS and pFG within each feature. Figure 5A summarized the results. Additionally, we performed the same analysis to compare the feature representation across 2 (feature: F_Emo_ and F_Id_) x 2 (node: ITG vs. MTG and STG).

For participant-wise source-level analyses, we again took the maximum MI across the 0–600 ms interval for each source and compared the lateral and ventral source distributions within each participant using independent-samples *t*-tests. Resulting p values were FDR-corrected across participants using FDR.

#### Task-relevant Feature Communications

We mapped feature-specific communication along the OCC–pSTS–TC for F_Emo_ and the OCC–pFG–TC for F_Id_. Sender and receiver sources were selected as those showing the maximum MI for the relevant feature: maximum F_Emo_ MI defined the OCC–pSTS and pSTS–TC pathways, whereas maximum F_Id_ MI defined the OCC-–pFG and pFG–TC pathways. Directed information (DI) was computed using event-related transfer entropy over a 0–600 ms post-stimulus window for the receiver, with a maximum sender delay of 400 ms. To isolate feature-specific communication, we computed DI conditioned on the feature (DI|F). Directed feature information (DFI) was defined as DI − DI|F, indexing communication information uniquely attributable to the feature. Significance was assessed with 200 permutation tests, using the 95th percentile of the null distribution as the threshold. DFI matrices were averaged across participants (F_Emo_: Emotion and Identity & Emotion; F_Id_: Identity and Identity & Emotion), and onset times of significant communication were extracted.

### Stage 3: Integration of F_Id_ and F_Emo_ in Temporal Cortex

#### Temporal Divergence of Known vs. Unknown identities

To compare F_Id_ representations between Known vs. Unknown identity, we analyzed task-relevant F_Id_ by computing MI(known F_Id_; MEG) and MI (unknown F_Id_; MEG) from 0 to 600 ms post animation onset, using the same statistical procedure as *Task-dependent Dynamic Feature Representations*. We examined MI for known and unknown separately in OCC, pFG, and TC. For each participant in Identity and Identity & Emotion groups, and for each source and condition (Known vs. Unknown), we extracted the maximum MI across time. We then averaged these values across sources within each ROI to yield one participant-level estimate for Known and Unknown F_Id_ in each region. Participant-level data were analyzed using a 2 (Known vs. Unknown) × 3 (OCC vs. pFG vs. TC) repeated-measures ANOVA with within-subject factors, followed by planned paired *t*-tests comparing known versus unknown identities within each region, with *p* values corrected across regions. Figure 6B and C show the corresponding results.

At the individual level, we again computed the maximum MI across time for each source and compared Known vs. Unknown F_Id_ representations across sources within each ROI using paired t-tests separately for each participant. The resulting participant-wise *p* values were corrected across participants within each ROI using false discovery rate (FDR) correction.

#### Feature Integration

As described in the main text (Figure 6), we quantified how TC integrates F_Id_ and F_Emo_ into a unified, person-relevant representation using information-theoretic synergy. Analyses focused on temporal regions where features from ventral and lateral pathways converge during the Identity & Emotion task.

##### Computation of Synergy

For each participant and each of the 792 sources within TC (ITG, MTG and STG), we computed Synergy between F_Id_, F_Emo_ and the MEG activity every 4 ms from 0–600 ms post animation onset:

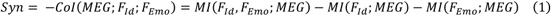

This measure quantifies the joint information between F_Id_ and F_Emo_ that exceeds the sum of their independent contributions to MEG responses. Higher values therefore index true feature integration rather than their addition. For each participant, this produced a 792-source × 150-time-point Synergy matrix describing the temporal evolution of feature interactions in TC.

##### Temporal Alignment of Integration

For each participant, we identified the source exhibiting maximum Synergy within the 0–600 ms window and extracted its time course. We also computed region-mean MI time courses for F_Id_ and F_Emo_ in TC. To relate the timing of integration of F_Id_ and F_Emo_ to feature availability in TC, we determined the later of the two MI peaks (F_Id_ and F_Emo_) across these regions, set that time to 0 ms, and realigned each participant’s maximal Synergy trace to this anchor. The aligned trace was visualized over a - 100 to +100 ms window to test whether Synergy increased after both features became locally represented.

##### Model-Based Verification of Integration

To confirm that the observed Synergy reflected genuine feature integration rather than their addition, we fitted explicit linear models to MEG activity of the source exhibiting maximum Synergy. For each participant, we modelled activity at time points of minimum and maximum Synergy at this source as a function of categorical factors F_Id_ and F_Emo_.

The baseline model included only main effects:

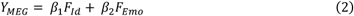

and the full model additionally included their interaction term,

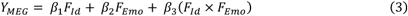

Both models were fitted using ordinary least squares. Model fit was quantified by the coefficient of determination R^2^, and the contribution of the interaction term was assessed using the log-likelihood ratio (LLR) between the full and baseline models:

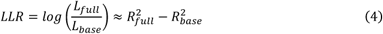

We compared LLR values between minimum and maximum Synergy using a paired t-test at the group level.

To generalize this effect from a single source to broader temporal regions, we repeated the analyses within TC. For each participant and source, we modelled activity at the time points of minimum and maximum mean Synergy (averaged across TC) and compared the corresponding LLR values. Group-level effects were tested using paired t-tests on the mean LLR across sources, and participant-level effects were assessed using paired t-tests across sources within each participant.

## Acknowledgements

This work was funded by the Wellcome Trust (Senior Investigator Award, UK; 107802) and the Multidisciplinary University Research Initiative/Engineering and Physical Sciences Research Council (USA, UK; 172046-01), awarded to Philippe G. Schyns; ERC [FACESYNTAX; 759796], awarded to Rachael E. Jack; the Wellcome Trust [214120/Z/18/Z], awarded to Robin A.A. Ince. The funders had no role in study design, data collection and analysis, decision to publish or preparation of the manuscript.

## Author Contributions

Conceptualization: PGS and YY. Experiment design: PGS and YY. Data curation: YY. Formal analysis: YY, PGS and RAAI. Funding acquisition: PGS and REJ. Resources: RAAI, OGBG, REJ and PGS. Software: OGBG and PGS. Visualisation: YY and PGS. Writing—original draft: YY and PGS. Writing—review and editing: all authors.

## Data and materials availability

Raw data and analyzed data have been deposited in Mendeley Data (https://doi.org/10.17632/dr5s3bgr82.1)

## References

1. Zhan, J., Liu, M., Garrod, O.G., Daube, C., Ince, R.A., Jack, R.E., and Schyns, P.G. (2021). Modeling individual preferences reveals that face beauty is not universally perceived across cultures. Current Biology 31, 2243–2252.

2. Hummert, M.L. (2014). Age changes in facial morphology, emotional communication, and age stereotyping. The Oxford Handbook of Emotion, Social Cognition, and Problem Solving in Adulthood, 47.

3. Todorov, A., Baron, S.G., and Oosterhof, N.N. (2008). Evaluating face trustworthiness: a model based approach. Social cognitive and affective neuroscience 3, 119–127.

4. Oosterhof, N.N., and Todorov, A. (2008). The functional basis of face evaluation. Proceedings of the National Academy of Sciences 105, 11087–11092.

5. Wolffhechel, K., Fagertun, J., Jacobsen, U.P., Majewski, W., Hemmingsen, A.S., Larsen, C.L., Lorentzen, S.K., and Jarmer, H. (2014). Interpretation of appearance: The effect of facial features on first impressions and personality. PloS one 9, e107721.

6. Bjornsdottir, R.T., Hensel, L.B., Zhan, J., Garrod, O.G., Schyns, P.G., and Jack, R.E. (2024). Social class perception is driven by stereotype-related facial features. Journal of Experimental Psychology: General.

7. Ekman, P., Sorenson, E.R., and Friesen, W.V. (1969). Pan-cultural elements in facial displays of emotion. Science 164, 86–88.

8. Rothbart, M., and Birrell, P. (1977). Attitude and the perception of faces. Journal of Research in Personality 11, 209–215.

9. van Rijsbergen, N., Jaworska, K., Rousselet, G.A., and Schyns, P.G. (2014). With Age Comes Representational Wisdom in Social Signals. Current Biology 24, 2792–2796. 10.1016/j.cub.2014.09.075.

10. Popham, S.F., Huth, A.G., Bilenko, N.Y., Deniz, F., Gao, J.S., Nunez-Elizalde, A.O., and Gallant, J.L. (2021). Visual and linguistic semantic representations are aligned at the border of human visual cortex. Nature Neuroscience 24, 1628–1636. 10.1038/s41593-021-00921-6.

11. Huth, A.G., Nishimoto, S., Vu, A.T., and Gallant, J.L. (2012). A continuous semantic space describes the representation of thousands of object and action categories across the human brain. Neuron 76, 1210–1224.

12. Ralph, M.A.L., Jefferies, E., Patterson, K., and Rogers, T.T. (2017). The neural and computational bases of semantic cognition. Nature reviews neuroscience 18, 42–55.

13. Nestor, A., Plaut, D.C., and Behrmann, M. (2011). Unraveling the distributed neural code of facial identity through spatiotemporal pattern analysis. Proceedings of the National Academy of Sciences 108, 9998–10003. 10.1073/pnas.1102433108.

14. Chang, L., and Tsao, D.Y. (2017). The Code for Facial Identity in the Primate Brain. Cell 169, 1013–1028.e14. 10.1016/j.cell.2017.05.011.

15. Zhan, J., Garrod, O.G., van Rijsbergen, N., and Schyns, P.G. (2019). Modelling face memory reveals task-generalizable representations. Nature human behaviour 3, 817–826.

16. O’Toole, A.J., Roark, D.A., and Abdi, H. (2002). Recognizing moving faces: a psychological and neural synthesis. Trends in Cognitive Sciences 6, 261–266. 10.1016/S1364-6613(02)01908-3.

17. Jack, R.E., Garrod, O.G., Yu, H., Caldara, R., and Schyns, P.G. (2012). Facial expressions of emotion are not culturally universal. Proceedings of the National Academy of Sciences 109, 7241–7244.

18. Jack, R.E., Garrod, O.G.B., and Schyns, P.G. (2014). Dynamic Facial Expressions of Emotion Transmit an Evolving Hierarchy of Signals over Time. Current Biology 24, 187–192. 10.1016/j.cub.2013.11.064.

19. Wong, S.S. (2016). Emotions and the communication of intentions in face-to-face diplomacy. European Journal of International Relations 22, 144–167.

20. Liu, M., Duan, Y., Ince, R.A., Chen, C., Garrod, O.G., Schyns, P.G., and Jack, R.E. (2022). Facial expressions elicit multiplexed perceptions of emotion categories and dimensions. Current Biology 32, 200–209.

21. Bracci, S., and Beeck, H.P.O. de (2023). Understanding Human Object Vision: A Picture Is Worth a Thousand Representations. Annual Review of Psychology 74, 113–135. 10.1146/annurev-psych-032720-041031.

22. Kanwisher, N., McDermott, J., and Chun, M.M. (1997). The Fusiform Face Area: A Module in Human Extrastriate Cortex Specialized for Face Perception. J. Neurosci. 17, 4302–4311. 10.1523/JNEUROSCI.17-11-04302.1997.

23. Grill-Spector, K., and Weiner, K.S. (2014). The functional architecture of the ventral temporal cortex and its role in categorization. Nature Reviews Neuroscience 15, 536–548.

24. Sliwinska, M.W., Bearpark, C., Corkhill, J., McPhillips, A., and Pitcher, D. (2020). Dissociable pathways for moving and static face perception begin in early visual cortex: Evidence from an acquired prosopagnosic. Cortex 130, 327–339.

25. Gao, X., Vuong, Q.C., and Rossion, B. (2019). The cortical face network of the prosopagnosic patient PS with fast periodic stimulation in fMRI. Cortex 119, 528–542.

26. Rezlescu, C., Pitcher, D., and Duchaine, B. (2012). Acquired prosopagnosia with spared within-class object recognition but impaired recognition of degraded basic-level objects. Cognitive Neuropsychology 29, 325–347.

27. Steeves, J.K., Culham, J.C., Duchaine, B.C., Pratesi, C.C., Valyear, K.F., Schindler, I., Humphrey, G.K., Milner, A.D., and Goodale, M.A. (2006). The fusiform face area is not sufficient for face recognition: evidence from a patient with dense prosopagnosia and no occipital face area. Neuropsychologia 44, 594–609.

28. Pitcher, D., Pilkington, A., Rauth, L., Baker, C., Kravitz, D.J., and Ungerleider, L.G. (2020). The human posterior superior temporal sulcus samples visual space differently from other face-selective regions. Cerebral Cortex 30, 778–785.

29. Yan, Y., Zhan, J., Garrod, O.G., Ince, R.A., Jack, R.E., and Schyns, P.G. (2025). The brain computes dynamic facial movements for emotion categorization using a third pathway. Proceedings of the National Academy of Sciences 122, e2423560122.

30. Pitcher, D. (2025). Neuropsychological evidence of a third visual pathway specialized for social perception. Nat Commun 16, 5774. 10.1038/s41467-025-61396-8.

31. Pitcher, D., and Ungerleider, L.G. (2021). Evidence for a third visual pathway specialized for social perception. Trends in Cognitive Sciences 25, 100–110.

32. Puce, A. (2024). From Motion to Emotion: Visual Pathways and Potential Interconnections. Journal of Cognitive Neuroscience, 1–24.

33. Duan, Y., Zhan, J., Gross, J., Ince, R.A., and Schyns, P.G. (2024). Pre-frontal cortex guides dimension-reducing transformations in the occipito-ventral pathway for categorization behaviors. Current Biology 34, 3392–3404.

34. Zhan, J., Ince, R.A., Van Rijsbergen, N., and Schyns, P.G. (2019). Dynamic construction of reduced representations in the brain for perceptual decision behavior. Current Biology 29, 319–326. e4.

35. Yu, H., Garrod, O.G., and Schyns, P.G. (2012). Perception-driven facial expression synthesis. Computers & Graphics 36, 152–162.

36. Ince, R.A., Giordano, B.L., Kayser, C., Rousselet, G.A., Gross, J., and Schyns, P.G. (2017). A statistical framework for neuroimaging data analysis based on mutual information estimated via a gaussian copula. Human brain mapping 38, 1541–1573.

37. Ince, R.A.A., van Rijsbergen, N.J., Thut, G., Rousselet, G.A., Gross, J., Panzeri, S., and Schyns, P.G. (2015). Tracing the Flow of Perceptual Features in an Algorithmic Brain Network. Scientific Reports 5, 17681. 10.1038/srep17681%2520 https://www.nature.com/articles/srep17681%2523supplementary-information.

38. Marr, D. (2010). Vision: A Computational Investigation into the Human Representation and Processing of Visual Information (MIT Press).

39. Yovel, G., and Belin, P. (2013). A unified coding strategy for processing faces and voices. Trends in Cognitive Sciences 17, 263–271. 10.1016/j.tics.2013.04.004.

40. Burton, A.M., Jenkins, R., and Schweinberger, S.R. (2011). Mental representations of familiar faces. British Journal of Psychology 102, 943–958. 10.1111/j.2044-8295.2011.02039.x.

41. Ekman, P., and Friesen, W.V. (1978). Facial action coding system. Environmental Psychology & Nonverbal Behavior.

42. Ince, R.A., Paton, A.T., Kay, J.W., and Schyns, P.G. (2021). Bayesian inference of population prevalence. Elife 10, e62461.

43. Bruce, V., and Young, A. (1986). Understanding face recognition. British journal of psychology 77, 305–327.

44. Haxby, J.V., Hoffman, E.A., and Gobbini, M.I. (2000). The distributed human neural system for face perception. Trends in cognitive sciences 4, 223–233.

45. Monroy, M., Cowen, A.S., and Keltner, D. (2022). Intersectionality in emotion signaling and recognition: The influence of gender, ethnicity, and social class. Emotion 22, 1980–1988. 10.1037/emo0001082.

46. Gill, D., Garrod, O.G.B., Jack, R.E., and Schyns, P.G. (2014). Facial movements strategically camouflage involuntary social signals of face morphology. Psychol Sci 25, 1079–1086. 10.1177/0956797614522274.

47. Kay, K.N., Weiner, K.S., and Grill-Spector, K. (2015). Attention reduces spatial uncertainty in human ventral temporal cortex. Current Biology 25, 595–600.

48. Yan, Y., Zhan, J., Ince, R.A., and Schyns, P.G. (2023). Network communications flexibly predict visual contents that enhance representations for faster visual categorization. Journal of Neuroscience 43, 5391–5405.

49. Yan, Y., Zhan, J., Garrod, O., Cui, X., Ince, R.A., and Schyns, P.G. (2023). Strength of predicted information content in the brain biases decision behavior. Current Biology 33, 5505–5514.

50. Liu, T.T., Fu, J.Z., Chai, Y., Japee, S., Chen, G., Ungerleider, L.G., and Merriam, E.P. (2022). Layer-specific, retinotopically-diffuse modulation in human visual cortex in response to viewing emotionally expressive faces. Nature Communications 13, 6302.

51. Timme, N., Alford, W., Flecker, B., and Beggs, J.M. (2014). Synergy, redundancy, and multivariate information measures: an experimentalist’s perspective. Journal of computational neuroscience 36, 119–140.

52. Delis, I., Ince, R.A.A., Sajda, P., and Wang, Q. (2022). Neural Encoding of Active Multi-Sensing Enhances Perceptual Decision-Making via a Synergistic Cross-Modal Interaction. J. Neurosci. 42, 2344. 10.1523/JNEUROSCI.0861-21.2022.

53. Schyns, P.G., Snoek, L., and Daube, C. (2022). Degrees of algorithmic equivalence between the brain and its DNN models. Trends in Cognitive Sciences.

54. Daube, C., Xu, T., Zhan, J., Webb, A., Ince, R.A., Garrod, O.G., and Schyns, P.G. (2021). Grounding deep neural network predictions of human categorization behavior in understandable functional features: The case of face identity. Patterns, 100348.

55. Zhou, C., Miao, M.-C., Chen, X.-R., Hu, Y.-F., Chang, Q., Yan, M.-Y., and Kuai, S.-G. (2022). Human-behaviour-based social locomotion model improves the humanization of social robots. Nature Machine Intelligence 4, 1040–1052. 10.1038/s42256-022-00542-z.

56. Gramfort, A., Luessi, M., Larson, E., Engemann, D., Strohmeier, D., Brodbeck, C., Goj, R., Jas, M., Brooks, T., Parkkonen, L., et al. (2013). MEG and EEG data analysis with MNE-Python. Frontiers in Neuroscience 7.

57. Taulu, S., and Simola, J. (2006). Spatiotemporal signal space separation method for rejecting nearby interference in MEG measurements. Physics in Medicine & Biology 51, 1759.

58. Benjamini, Y., and Hochberg, Y. (1995). Controlling the False Discovery Rate: A Practical and Powerful Approach to Multiple Testing. Journal of the Royal Statistical Society: Series B (Methodological) 57, 289–300. 10.1111/j.2517-6161.1995.tb02031.x.

