## Supplemental Information for "Task demands dynamically structure feature selection, routing, and integration in the human brain"

### Supplementary Information

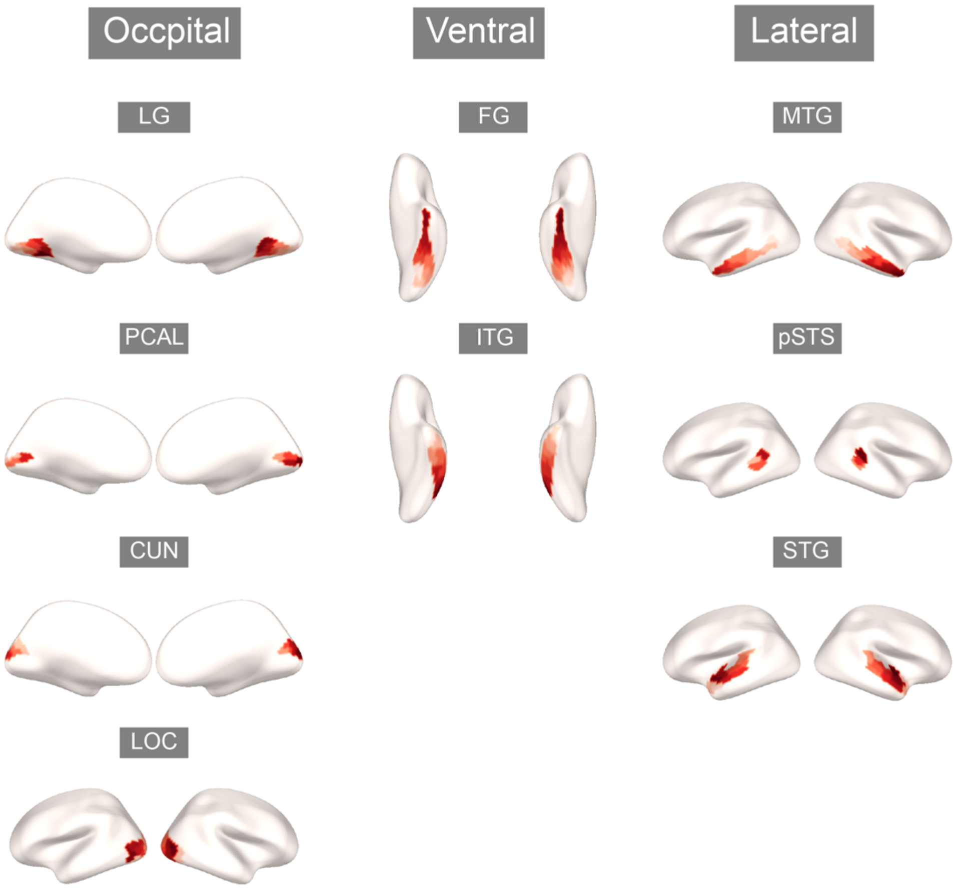


**Figure S1. Subregions forming the occipito-dorsal, ventral and social pathways.** Each inflated brain highlights a labelled region within OCC, ventral and lateral pathways, with constituent subregions color-coded.

**Table S1. Occipital representational persistence at the individual-level**

| | Participants | $F_{Id}$ last sig. time (ms) | $F_{Emo}$ last sig. time (ms) | Difference (ms) | *t* | $p_{FDR}$ |
|---|---|---|---|---|---|---|
| Emotion Task | 1 | 214 | 443 | -229 | -40.54 | < .001 |
| | 2 | 3 | 397 | -394 | -49.81 | < .001 |
| | 3 | 147 | 506 | -359 | -58.01 | < .001 |
| | 4 | 1 | 541 | -540 | -144.44 | < .001 |
| | 5 | 1 | 475 | -474 | -71.51 | < .001 |
| | 6 | 352 | 581 | -229 | -34.09 | < .001 |
| | 7 | 8 | 376 | -368 | -43.77 | < .001 |
| | 8 | 310 | 589 | -279 | -53.01 | < .001 |
| Identity Task | 9 | 477 | 60 | 416 | 58.81 | < .001 |
| | 10 | 301 | 1 | 300 | 37.25 | < .001 |
| | 11 | 468 | 503 | -35 | -5.11 | < .001 |
| | 12 | 548 | 269 | 280 | 34.08 | < .001 |
| | 13 | 539 | 3 | 536 | 138.83 | < .001 |
| | 14 | 562 | 404 | 158 | 26.51 | < .001 |
| | 15 | 526 | 209 | 317 | 48.61 | < .001 |
| | 16 | 536 | 269 | 267 | 36.62 | < .001 |

**Table S2. Selective routing of $F_{Emo}$**

| | Participants | Mean MI in pSTS (bits) | Mean MI in pFG (bits) | Difference (bits) | *t* | $p_{FDR}$ |
|---|---|---|---|---|---|---|
| Emotion Task | 1 | 0.0012 | 0.0092 | 0.0023 | 4.08 | < .001 |
| | 2 | 0.0096 | 0.0039 | 0.0056 | 12.13 | < .001 |
| | 3 | 0.014 | 0.0078 | 0.00063 | 8.77 | < .001 |
| | 4 | 0.013 | 0.0068 | 0.0057 | 19.00 | < .001 |
| | 5 | 0.013 | 0.0052 | 0.0081 | 19.98 | < .001 |
| | 6 | 0.012 | 0.0012 | -0.00030 | -0.59 | .556 |
| | 7 | 0.011 | 0.0050 | 0.0064 | 15.30 | < .001 |
| | 8 | 0.019 | 0.012 | 0.0063 | 8.52 | < .001 |
| Identity & Emotion Task | 17 | 0.013 | 0.0059 | 0.0074 | 21.60 | < .001 |
| | 18 | 0.011 | 0.0060 | 0.0053 | 20.61 | < .001 |
| | 19 | 0.0013 | 0.0001 | 0.0012 | 3.08 | < .001 |
| | 20 | 0.011 | 0.0062 | 0.0047 | 16.44 | < .001 |
| | 21 | 0.0073 | 0.0026 | 0.0047 | 8.74 | < .001 |
| | 22 | 0.012 | 0.0010 | 0.0023 | 4.83 | < .001 |
| | 23 | 0.011 | 0.0080 | 0.0032 | 9.37 | < .001 |
| | 24 | 0.013 | 0.0078 | 0.0046 | 11.20 | < .001 |

**Table S3. Selective routing of $F_{Id}$**

| | Participants | Mean MI in pSTS (bits) | Mean MI in pFG (bits) | Difference (bits) | *t* | $p_{FDR}$ |
|---|---|---|---|---|---|---|
| Identity Task | 9 | 0.0072 | 0.010 | -0.0031 | -5.92 | < .001 |
| | 10 | 0.0066 | 0.0089 | -0.0023 | -3.89 | < .001 |
| | 11 | 0.0063 | 0.0083 | -0.0020 | -3.36 | .0013 |
| | 12 | 0.0067 | 0.0094 | -0.0027 | -5.58 | < .001 |
| | 13 | 0.019 | 0.017 | 0.0020 | 1.42 | .17 |
| | 14 | 0.009 | 0.0012 | -0.0028 | -6.27 | < .001 |
| | 15 | 0.0069 | 0.0076 | -0.00067 | -2.19 | .036 |

| | | | | | | |
|---|---|---|---|---|---|---|
| | 16 | 0.0090 | 0.0012 | -0.0028 | -6.27 | < .001 |
| Identity & Emotion Task | 17 | 0.0081 | 0.016 | -0.0079 | -6.54 | < .001 |
| | 18 | 0.0011 | 0.0011 | 0 | -0.11 | .91 |
| | 19 | 0.00022 | 0.0022 | -0.0020 | -6.14 | < .001 |
| | 20 | 0.0028 | 0.0046 | -0.0018 | -3.59 | < .001 |
| | 21 | 0.0032 | 0.0015 | 0.0017 | 3.70 | < .001 |
| | 22 | 0.0066 | 0.0098 | -0.0031 | -4.81 | < .001 |
| | 23 | 0.0012 | 0.0036 | -0.0024 | -5.46 | < .001 |
| | 24 | 0.0037 | 0.0047 | -0.0011 | -2.00 | .054 |

**Table S4 Feature ($F_{Emo}$ vs. $F_{Id}$) × Region (ITG vs. MTG/STG) ANOVA (TC)**

| | *F* | Num DF | Den DF | *p* |
|---|---|---|---|---|
| Feature | 0.060 | 1 | 15 | .81 |
| Region | 1.17 | 1 | 15 | .30 |
| Feature ×Region | 1.99 | 1 | 15 | .18 |

**Table S5. Divergence of Known vs. Unknown $F_{Id}$ (pFG)**

| | Participants | Mean MI for Known $F_{Id}$ (bits) | Mean MI for Unknown $F_{Id}$ (bits) | Difference (bits) | *t* | $p_{FDR}$ |
|---|---|---|---|---|---|---|
| Identity Task | 9 | 0.016 | 0.0012 | 0.015 | 21.88 | < .001 |
| | 10 | 0.0039 | 0.0052 | -0.0013 | -1.78 | .095 |
| | 11 | 0.0079 | 0.0014 | 0.0066 | 11.55 | < .001 |
| | 12 | 0.011 | 0.0068 | 0.0046 | 7.37 | < .001 |
| | 13 | 0.013 | 0.0082 | 0.0045 | 7.54 | < .001 |
| | 14 | 0.011 | 0.0056 | 0.0059 | 10.42 | < .001 |
| | 15 | 0.012 | 0.010 | 0.0017 | 2.48 | .021 |
| | 16 | 0.0069 | 0.0022 | 0.0046 | 9.21 | < .001 |
| Identity & Emotion Task | 17 | 0.016 | 0.014 | 0.0017 | 2.89 | .0071 |
| | 18 | 0.00026 | 0.00007 | 0.00019 | 1.13 | .30 |
| | 19 | 0.0015 | 0.00071 | 0.00075 | 1.838 | .091 |
| | 20 | 0.0019 | 0.00018 | 0.00012 | 0.22 | .83 |
| | 21 | 0.0004 | 0.00019 | 0.00021 | 0.77 | .472 |
| | 22 | 0.010 | 0.0061 | 0.0042 | 6.62 | < .001 |
| | 23 | 0.0001 | 0.0021 | -0.0020 | -5.03 | < .001 |
| | 24 | 0.0037 | 0.00028 | 0.0034 | 5.45 | < .001 |

**Table S6. Divergence of Known vs. Unknown $F_{Id}$ (TC)**

| | Participants | Mean MI for Known $F_{Id}$ (bits) | Mean MI for Unknown $F_{Id}$ (bits) | Difference (bits) | *t* | $p_{FDR}$ |
|---|---|---|---|---|---|---|
| Identity Task | 9 | 0.0073 | 0 | 0.0073 | 30.24 | < .001 |
| | 10 | 0.0012 | 0.00032 | 0.00087 | 5.86 | < .001 |
| | 11 | 0.0048 | 0.00044 | 0.0044 | 20.99 | < .001 |
| | 12 | 0.0068 | 0.0027 | 0.0041 | 16.02 | < .001 |
| | 13 | 0.0074 | 0.0066 | 0.00083 | 2.84 | < .001 |
| | 14 | 0.0069 | 0.0013 | 0.0056 | 27.70 | < .001 |
| | 15 | 0.0044 | 0.0016 | 0.0028 | 12.25 | < .001 |
| | 16 | 0.0055 | 0.00038 | 0.0051 | 25.22 | < .001 |
| | 17 | 0.0082 | 0.0059 | 0.0023 | 10.62 | .0071 |

| | | | | | | |
|---|---|---|---|---|---|---|
| Identity & Emotion Task | 18 | 0.00076 | 0.00060 | 0.00016 | 1.09 | .27 |
| | 19 | 0.00039 | 0.00008 | 0.00031 | 3.76 | < .001 |
| | 20 | 0.0010 | 0.0039 | 0.00061 | 4.28 | < .001 |
| | 21 | 0.00075 | 0.00016 | 0.00059 | 4.86 | < .001 |
| | 22 | 0.0045 | 0.0038 | 0.00075 | 3.06 | .0026 |
| | 23 | 0.0009 | 0.00057 | -0.00047 | -4.43 | < .001 |
| | 24 | 0.0029 | 0.00005 | 0.0029 | 13.58 | < .001 |
